# Limited bone marrow chimerism impairs cell competition in the thymus and causes leukemia

**DOI:** 10.64898/2025.12.16.694652

**Authors:** Bruna S. P. Oliveira, Rafael A. Paiva, Carolina C. Pinto, Rita V. Paiva, Michelle X. G. Pereira, Camila V. Ramos, Sara Azenha, Francisca Xara-Brasil, Alexandre Kaizeler, Tiago Paixão, Pedro Faísca, Vera C. Martins

## Abstract

Cell competition in the thymus is a critical tumor suppressor process that prevents leukemia. We show that suboptimal bone marrow correction of _γ*c*_-deficient mice disrupts this process, inducing thymus autonomy and promoting T cell acute lymphoblastic leukemia (T-ALL). _γ*c*_-deficiency causes severe combined immunodeficiency in mice and humans and is treated by hematopoietic stem and progenitor cell (HSPC) transplantation. However, inefficient correction led to intermittent thymic activity, consistent with sporadic thymus seeding. This was accompanied by altered thymocyte composition and the emergence of an aberrant TCRβ□CD4□CD8□ population that preceded overt leukemia. Leukemia incidence and latency were exacerbated when HSPCs were pre-cultured under conditions mimicking gene-editing protocols, whereas delaying thymopoietic support in the host mitigated disease. Critically, the extent of bone marrow reconstitution determined the ability to suppress thymus autonomy and leukemia development. Together, these findings demonstrate that inadequate bone marrow correction directly compromises thymic cell competition, creating a permissive niche for malignant transformation. Our work highlights thymus autonomy as a risk in immunodeficiency treatment and underscores the need to ensure robust hematopoietic reconstitution to prevent leukemogenesis.

## INTRODUCTION

Cell competition in the thymus is a process that promotes cellular turnover and thereby suppresses leukemia^1^. Cell competition was originally described in *Drosophila*^2^, and considers that cell clones of different fitness co-occur in a tissue and the fittest outcompete less fit cells, thereby contributing to optimal organ function^3^. Cell competition is a fundamental process in embryonic development, hematopoiesis, cardiomyocyte homeostasis in mice, and liver regeneration in rats^4^. It counteracts the pathogenesis of intestinal cancer by promoting the extrusion of transformed epithelial cells into the intestinal lumen^5^, and regulates cell growth in human cancer^6^. In the thymus, cell competition favors the turnover and replenishment of a precursor population, thereby providing a constant pool of optimal cells for T lymphocyte differentiation, and this suppresses leukemia^1^.

Using an experimental mouse model of thymus transplantation, we showed that young T cell precursors, which recently seeded the thymus, are essential to purge the old, less fit precursors, with longer thymus residency^1^. This occurs at defined stages of T cell differentiation and is not generalized to all thymocytes^7^. If cell competition in the thymus is impaired or abrogated, i.e., if thymus colonization by proficient bone marrow progenitors is blocked, the thymus autonomously sustains T lymphocyte production and export for months – thymus autonomy^8, 9^. But there is a major trade-off: thymus autonomy leads to malignant transformation of T lymphocyte precursors and the onset of T cell acute lymphoblastic leukemia (T-ALL)^1, 10^. The murine disease closely resembles human T-ALL in terms of immunophenotype, pathology, and genetic alterations^1^. Further similarities between the mouse and human T-ALL were gain-of-function mutations in *Notch1*^11^, as well as the expression of *Tal1* and *Lmo2* oncogenes, a molecular hallmark of the largest subgroup of pediatric T-ALL^12, 13^. T-ALL is a particularly aggressive subtype of acute lymphoblastic leukemia that arises from T lymphocyte precursors^14^, and accounts for 10-15% of pediatric and 20-25% of adult acute lymphoblastic leukemia (ALL) cases^15, 16^. In this context, we propose that leukemia emerging from impaired cell competition in the murine thymus follows a conserved process that informs on mechanisms underlying human T-ALL.

Our former results resembled the outcome of gene therapy to correct X-linked SCID-X1 in the early 2000’s^17, 18^. SCID-X1 is the most common subtype of severe combined immunodeficiency (SCID), and is caused by loss-of-function mutations in the *common gamma chain* (_γ*c*_) gene^19^. SCID-X1 is lethal if untreated because of the lack of T and NK cells, and the presence of non-functional B lymphocytes^20^, which render affected baby boys susceptible to recurrent infections and failure to thrive. While bone marrow transplantation is the standard of care^19, 21^, this approach is limited by the requirement of compatible donors. Being a monogenetic disease, SCID-X1 is a prime candidate for gene therapy treatment. Indeed, _γ*c*_ gene correction by gene therapy delivered the first successful results in the early 2000s, with thymus function effectively established and *de novo* T lymphocyte production^22, 23^. While this achievement speaks in favor of SCID-X1 being a good candidate for gene therapy treatment, there were caveats to this approach. Thymus function was kept active in the treated patients, despite indication that the bone marrow ceased to contribute to thymopoiesis^24^, a condition compatible with thymus autonomy. The lack of bone marrow contribution indicates that long-term reconstitution was not reached, or its levels were so low that remained undetected. This was expected because those patients were not myeloablated prior to treatment^22, 23^. After gene therapy, 25% of the treated patients developed T-ALL^18, 25^, which was attributed to genotoxicity caused by vector insertion next to oncogenes^26, 27^. We have proposed an alternative explanation: that thymus autonomy triggered leukemia in the gene therapy-treated patients, and that any condition favoring thymus autonomy is permissive to T-ALL. This notion has received support from two independent studies that showed that low level of _γ*c*_-proficient reconstitution in bone marrow chimeras has a high risk of leukemia, independently of gene therapy^28, 29^. SCID-X1 remains an unmet medical need that requires novel therapeutic approaches. This requires the discovery of the principles underlying bone marrow and thymus communication, and how this controls cell competition and T cell development. Adding to the remarkable efforts taken to develop safe viral vectors for gene therapy, we propose to dissect how the bone marrow correction of _γ*c*_-deficiency determines cell fate decisions in the thymus, i.e., whether normal thymopoiesis or leukemogenesis will be favored.

We show that non-preconditioned _γ*c*_-deficient mice develop T-ALL following transplantation of a reduced proportion of wild type hematopoietic stem and progenitor cells (HSPC). Preceding T-ALL, thymopoiesis exhibited features that were consistent with the disruption of cell competition, and an early stage of thymus autonomy: DN3-early were highly proliferative, had increased CD71 expression, and a population of CD4^+^CD8^+^ cells lacking TCRβ expression emerged. The latter population also emerged from *Rag2^-/-^* donors, that have a developmental block at the DN3-early, prior to giving rise to T-ALL. Finally, we found that the level of bone marrow reconstitution defines its capacity to inhibit thymus autonomy upon hematopoietic stem and progenitor cell (HSPC) transplantation. Collectively, our findings establish thymus autonomy as a thymus-intrinsic property with significant implications for health, underscoring the need to restrain it during manipulation of hematopoiesis, and bone marrow reconstitution in a setting of immunodeficiency.

## RESULTS

### Inefficient bone marrow reconstitution causes T cell acute lymphoblastic leukemia (T-ALL)

We have previously shown that thymus autonomy has a high risk of T-ALL^1, 10^, and proposed that every condition triggering thymus autonomy is permissive to T-ALL^1, 30^. To test whether T-ALL emerges from wild type hematopoietic progenitors used to reconstitute the bone marrow of _γ*c*_-deficient mouse hosts in limited proportions, we injected HSPCs intravenously in a mixture of 10% wild type and 90% _γ*c*_*^-/-^* into _γ*c*_*^-/-^* mice^31^ without preconditioning (Fig. 1 A). HSPCs were either immediately injected after isolation, or cultured for 16 hours using a cytokine cocktail that mimicked the conditions used for gene editing^29^ (Fig. 1 A). No genetic manipulation was performed. While T-ALL developed in both conditions, it had a faster onset and higher incidence in the group injected with pre-cultured HSPCs (Fig. 1 B). Specifically, mice injected with pre-cultured HSPCs developed T-ALL with an onset at 20.4 week-post injection and incidence of 75.0%, while mice injected with non-cultured HSPCs had the onset of T-ALL at week 36.9 and an incidence of 46.1% (Fig. 1 B). Despite the phenotypic variability, leukemias had an immature phenotype and the most common resembled CD8 immature single positive to double positive thymocytes (Fig. 1 C). The cellularity of thymi and spleens in T-ALL-bearing mice was overly increased as compared to that of healthy wild type controls (Fig. 1 D). Most organs analyzed were infiltrated with T-ALL blasts, with a diffuse neoplastic lymphoid proliferation with multisystemic involvement (Fig. 1 E, F). In the central nervous system, the infiltrates involved the arachnoid and pia mater, extending into the choroid plexus of the fourth ventricle. The bone marrow was completely replaced by a diffuse lymphoid neoplastic population. In the lungs, there was marked perivascular and peribronchiolar infiltration by neoplastic lymphoid cells. The spleen showed extensive lymphoid proliferation with complete obliteration of the normal splenic architecture. In the liver, lymphoid cells proliferated in a portal distribution, effacing the portal tracts and invading the sinusoidal capillaries. The kidneys exhibited interstitial, capsular, and hilar infiltration by neoplastic lymphoid cells, and similar infiltrates were observed in the subepicardium of the right ventricle and right atrium of the heart. In sum, T-ALL emerged from wild type HSPCs transplanted in restricted proportions into non-irradiated _γ*c*_*^-/-^* hosts without genetic manipulation. Pre-culturing HSPCs had an impact on disease dynamics by increasing incidence and accelerating the onset of T-ALL.

**Figure 1.**
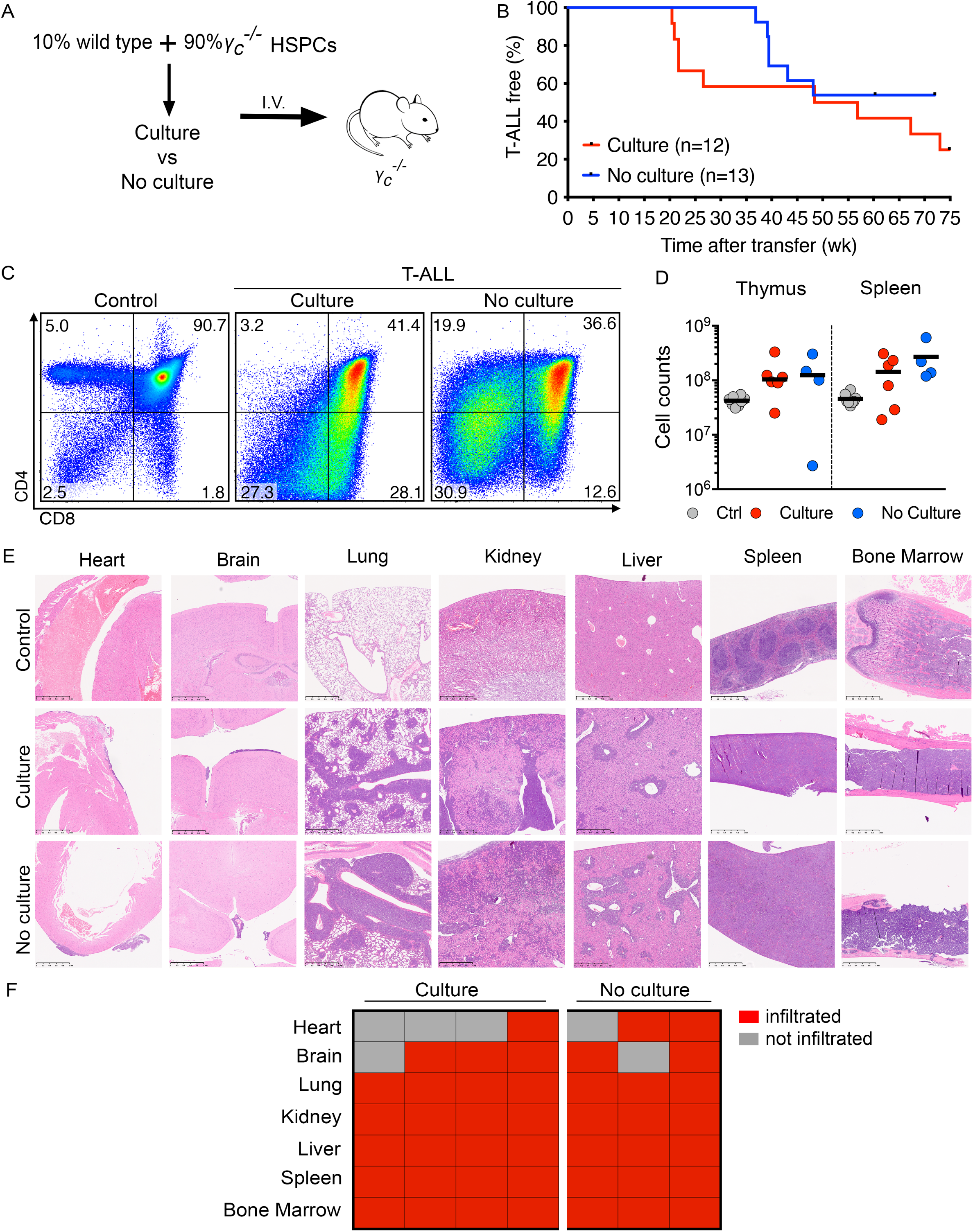
Limited bone marrow chimerism causes T-ALL. A) 10^6^ HSPCs in a mixture of 10% wild type plus 90% _γ*c*_*^-/-^*, were i.v. injected into non-preconditioned _γ*c*_*^-/-^* hosts. One group of mice received HSPCs that were previously cultured for 16 hours as indicated. CD45.1 and CD45.2 were used to distinguish donor/host-derived cells. B) Percentage of T-ALL free mice over time. C) Representative CD4/CD8 profiles of wild type control thymocytes and T-ALL samples. Numbers show percentage of cells in each quadrant, gated on live donor cells for T-ALL. D) Cellularity of thymi and spleens in healthy wild type controls and mice with T-ALL. Each dot represents one mouse, and the lines represent the medians. E) Representative sections of the indicated organs stained with hematoxylin and eosin. Scale bars: 500μm. F) Overview of mice shown in (E).

### Inefficient bone marrow reconstitution promotes inter-individual thymic heterogeneity with prevalence of DN3 as the most immature thymocytes

To test whether thymus autonomy preceded T-ALL, we used the same protocol as in Fig. 1A and analyzed the mice at 3-, 9-, and 15-week post-injection. There was considerable heterogeneity in the analyzed thymi among different mice. While thymopoiesis was active in most samples, a minority was composed only of circulating T cells (Fig. 2A, B), consistent with a past wave of thymopoiesis. Total thymus cellularity was highest at 3 weeks and declined thereafter (Fig. 2 C). The most immature CD4/CD8 donor double negative (DN) stages were also phenotypically heterogeneous. The presence of the most immature wild type HSPC-derived thymocyte populations, the early T lineage progenitor (ETP) and DN2 suggests recent thymus colonization and could be found only in a minority of the analyzed thymi (Fig. 2 D, E, F, Supplementary Fig.1). Of note, donor DN3 were detected in most thymi at all time points of analysis (Fig. 2 D, G) and were often the most immature thymocytes present (Fig. 2 D, right plot, and Supplementary Fig.1). This was, in fact, the most common phenotype at all timepoints, corresponding to 73.4%, 52.6%, or 52.6% of the analyzed thymi at 3-, 9- or 15-week post injection, respectively (Supplementary Fig.1). In physiology, the expression of the transferrin receptor CD71 is restricted to rare DN3-early cells, and is upregulated following β-selection, as thymocytes differentiate into DN3-late. This differs in thymus autonomy, where the level of CD71expression DN3-early is increased^32^. Here, CD71 expression was heterogeneous across samples, with some samples identical to healthy controls, and others with phenotypes similar to those described at early stages of thymus autonomy (Fig. 2 H, I, J)^32^. Likewise, Ki-67 expression (used as proxy for proliferation) was heterogeneous, with samples resembling healthy controls and others identical to autonomous thymi (Fig. 2 K). Cell numbers of DN3-early reduced progressively in time, but the population could be detected for at least 15 weeks (Fig. 2 L). Lastly, we searched for an abnormal CD4^+^CD8^+^ population lacking TCRβ, which preceded leukemia in a thymus transplantation model^32^. Indeed, we found a TCRβ-negative population within CD4^+^CD8^+^ cells (Fig. 2 M), that accumulated in several samples (Fig. 2 N, O). Several hematopoietic progenitors were analyzed in these mice (Supplementary Fig. 2A), and a low but detectable level of engraftment was found in LSK and common lymphoid progenitor (CLP, Supplementary Fig. 2B), for up to 15 weeks post injection. Albeit at extremely low levels, multipotent progenitors 1 (MPP1s, Supplementary Fig. 2C, D), as well as the myeloid-bias MPP2 and MPP3 (Supplementary Fig. 2C, D), also contained wild type donor derived cells. Altogether, these data confirm that bone marrow reconstitution by few injected HSPCs is extremely inefficient in non-irradiated _γ*c*_*^-/-^* hosts. Most thymi lacked the most immature thymocyte populations but DN3 remained, suggesting that thymopoiesis could be kept active as result of persistence of this population, as previously reported^32^.

**Figure 2.**
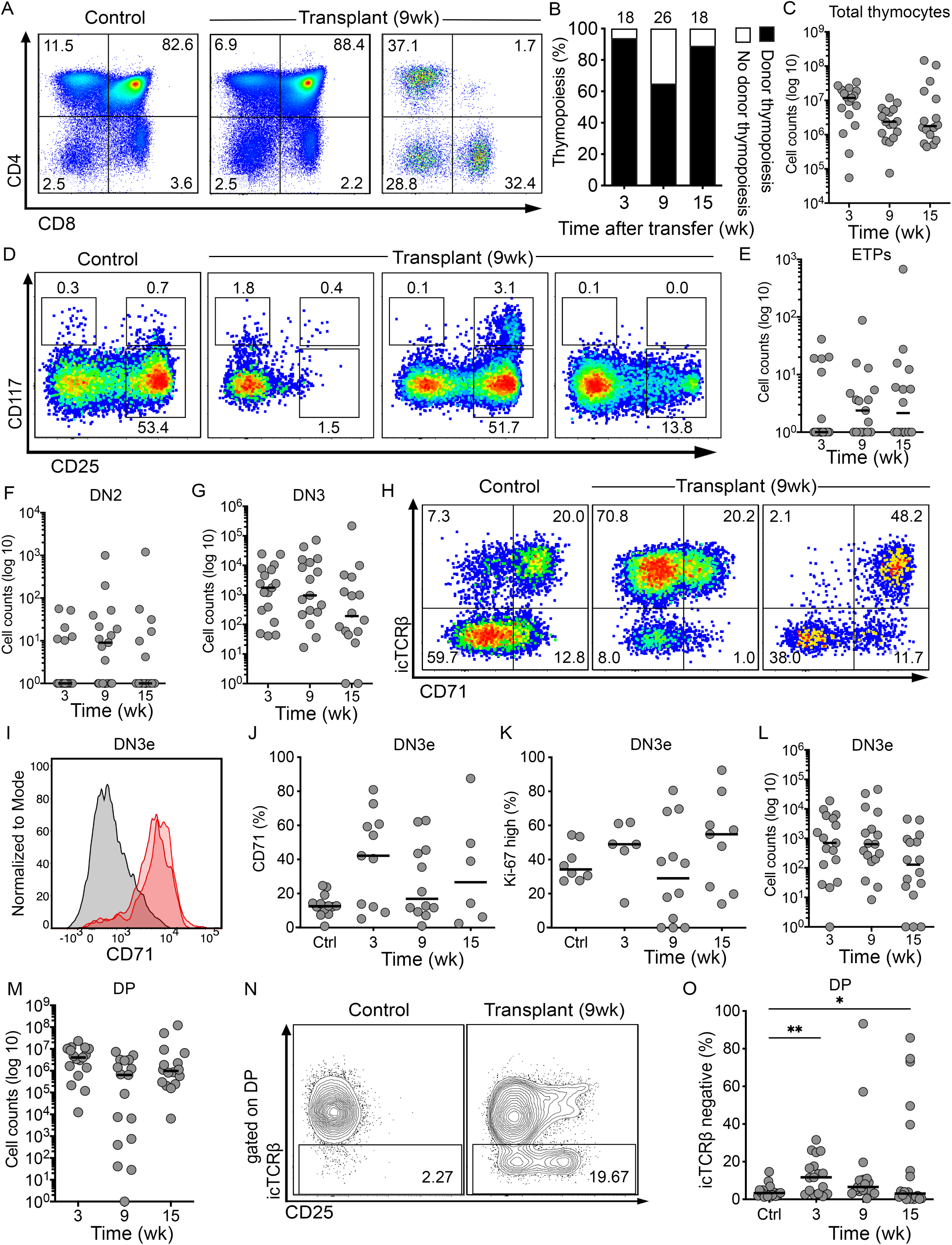
Low-level bone marrow chimerism associates with phenotypic heterogeneity and hallmarks of thymus autonomy. A) Representative examples of the two most extreme phenotypes obtained for the donor CD4/CD8 profile following bone marrow transplantation. Examples are from 9-week post-transplantation. Numbers show percentage of cells in each quadrant, gated on donor live thymocytes. B) Quantification of thymi with or without donor-derived thymopoiesis at 3-, 9- and 15-week post-transplantation, considered as those with at least 10% donor CD4/CD8 thymocytes. Numbers on top of the columns indicate the number of mice analyzed. C) Thymus cellularity at the indicated time points. D) Representative examples of the phenotypes observed among the most immature donor DN thymocyte populations. Examples are from 9 weeks after HSPC transplantation, and cells were pre-gated as negative for CD4, CD8, Lin-markers. E-G) Absolute cell counts of the indicated populations 3-, 9- and 15-week post-injection. H) Donor DN3 thymocytes are shown for the intracellular expression of TCRβ and CD71. I) Example of donor DN3-early samples 9-wk after injection (red) or a control (grey). J) Quantification of CD71+ cells in DN3-early (DN3e) cells. K) Quantification of Ki67-high donor DN3-early. L) Cellularity of donor DN3-early. M) Quantification of donor derived CD4^+^CD8^+^ double positive (DP) cells. N) Representative example of one control versus one thymus resulting from limited bone marrow chimerism, gated on CD4^+^CD8^+^ cells. O) Quantification of TCRβ-negative cells in CD4^+^CD8^+^ donor thymocytes. Each circle represents one mouse and the horizontal line is the median. Control samples refer to wild type, non-manipulated mice. Data are from 10 independent experiments containing a total n = 62.

### TCRβ^-^CD4^+^CD8^+^ aberrant cells develop from autonomous DN3-early thymocytes

As the former analyses did not permit to accurately distinguish thymus autonomy from discrete waves of thymopoiesis, we sought to minimize cellular complexity in the thymus and test whether DN3-early were sufficient to generate the aberrant TCRβ^-^CD4^+^CD8^+^ cells. We therefore used a mixture of 10% *Rag2^-/-^* (with or without prior culture) plus 90% _γ_*_c_^-/-^* HSPCs injected into _γ_*_c_^-/-^* mice, which were analyzed 9 or 15 weeks later (Fig. 3 A). Thymus cellularity was reduced from the first to the second time point (Fig. 3 B). In line with the previous results, *Rag2^-/-^* donor DN3 thymocytes were detected at both time points, including in thymi lacking the most immature ETP and DN2 populations (Fig. 3 C, D). TCRβ^-^CD4^+^CD8^+^ cells were detected in all analyzed thymi (Fig. 3 E, F). Not every thymus with CD4^+^CD8^+^ cells had a population of DN3 (Fig. 3 D, F). No differences in relative or absolute cell counts were detected between the thymi from cultured versus non-cultured *Rag2^-/-^* donors for DN3-early (Supplementary Fig. 3 A, B) and TCRβ^-^CD4^+^CD8^+^ cells (Supplementary Fig. 3 C, D). Consistent with the results obtained with wild type HSPC donors (Fig. 1), T-ALL also ensued from *Rag2^-/-^* HSPC, with a total of 9 T-ALLs detected from a total of 27 injected hosts in 4 independent experiments (not shown). Many of these mice died from causes unrelated to T-ALL, which precluded construction of a T-ALL-free survival curve. Nevertheless, our results show that *Rag*-deficient thymocytes can bypass the β-selection checkpoint in a scenario of limited HSPC reconstitution of _γ_*_c_^-/-^* hosts, generate TCRβ^-^CD4^+^CD8^+^ cells, and are sufficient to cause T-ALL. Next, to test our hypothesis that thymus seeding was intermittent, we reasoned that every progenitor immigrating into the thymus generates T cells that are exported to the blood, which can be monitored over time. To distinguish recent thymic emigrants (RTE), we used HSPCs from *Tg(Rag2p-GFP)* mice^27, 28, 33^, in which T cells maintain GFP expression for up to 10 days after thymus egress^34^. A mixture of 10% *Tg(Rag2p-GFP)* and 90% _γ_*_c_^-/-^* was injected into _γ_*_c_^-/-^* mice, and their blood was monitored thereafter (Fig. 3 G). Both B and T GFP-positive cells were detected in the blood at different, non-coincident, time points (Fig. 3 H). Importantly, clear peaks of GFP-positive T lymphocytes in the blood were interspersed by periods without detection of GFP-positive T cells (Fig. 3 I). These data demonstrate that T lymphocyte production was not constant overtime and could be jumpstarted even after long periods lacking thymic activity (Fig. 3 H, I, top panels). These refractory periods varied in duration across individual mice, suggesting stochasticity in thymus seeding and subsequent thymopoiesis. In line with the results shown in Fig. 1, these mice developed T-ALL with onset at 29.1 week-post injection and incidence of 66.7% by the end of the observation period (Fig. 3 J). The immunophenotype of these leukemias was heterogeneous, but commonly with an immature CD4+CD8+ profile (Fig. 3 K). Together, these data show that T lymphopoiesis resulting from low-level chimerism of the bone marrow is intermittent, which can be explained by a stochastic pattern of thymus colonization by rare wild type progenitors, that ultimately culminates in T-ALL.

**Figure 3.**
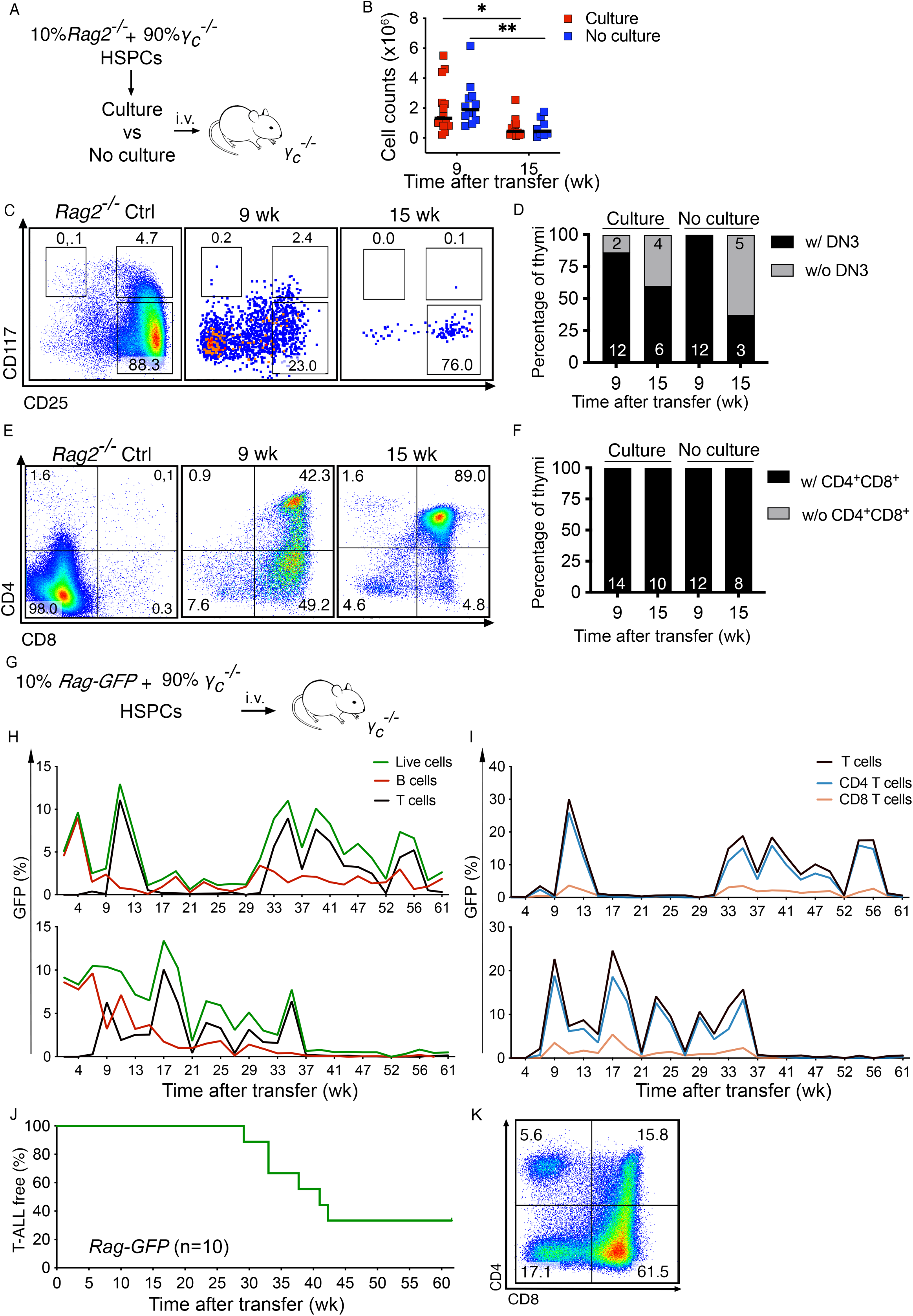
TCRβ^-^CD4^+^CD8^+^ cells and intermittent thymopoiesis precede T-ALL. A) 10^6^ HSPCs in a mixture of 10% *Rag2^-/-^* plus 90% _γ*c*_*^-/-^*, were i.v. injected into non-preconditioned _γ*c*_*^-/-^* mice, which were analyzed 9 and 15 weeks later. CD45.1 and CD45.2 were used to distinguish donor/host-derived cells. B) Thymus cellularity after HSPC transplantation. Each dot represents one mouse, and the lines represent the medians. C) Representative examples of donor DN populations. Cells were pre-gated as negative for CD4, CD8, Lin-markers. D) Quantification of thymi with or without DN3 thymocytes. E) Representative examples of thymi with a population of donor CD4+CD8+. F) Quantification of samples with a CD4+CD8+ double positive (DP-like) population. Data in E-F is cumulative from 8 independent experiments with n=4-8 mice/experiment and a total of n=20 without culture, and n=24 with culture. G) 10^6^ HSPCs in a mixture of 10% *Tg(Rag2-GFP)* plus 90% _γ*c*_*^-/-^*, were i.v. injected into non-preconditioned _γ*c*_*^-/-^* mice, which were periodically bled thereafter. H) Quantification of GFP-positive cells in the blood, as indicated. Each graph corresponds to one mouse. I) Quantification of GFP-positive cells in the blood, as percentage of total T cells, and the correspondent proportion of CD4+ and CD8+ T cells, over time. Each graph represents the same mice in H. J) Percentage of T-ALL-free over time. K) Representative CD4/CD8 FACS profile of one T-ALL. Numbers show percentage of cells in each quadrant. Data in A-F are from 8 independent experiments containing n =4-8/experiment. Data in H-K are from one individual experiment, n=10.

### T-ALL incidence and onset depend on the thymic microenvironment

Former studies using _γ*c*_-deficient^29^ or *Rag2^-/-^*_γ*c*_*^-/-^* mice^28^ reported some differences in T-ALL incidence, and we therefore injected HSPCs in a mixture of pre-cultured 10% wild type plus 90% _γ*c*_*^-/-^* into *Rag2^-/-^*_γ*c*_*^-/-^* recipients (Fig. 4 A). Although T-ALL also emerged in *Rag2^-/-^*_γ*c*_*^-/-^*hosts (Fig. 4 B), it had a later onset (47.9 wk) and lower incidence (22.2%) when compared to the _γ*c*_*^-/-^* recipients (Fig. 1 B). T-ALL blasts were immature single positive to CD4+CD8+ (Fig. 4 C), and the cellularity of both thymi and spleens was overly enlarged as compared to a wild type healthy control (Fig. 4 D). All analyzed organs were infiltrated with blasts in T-ALL-bearing mice (Fig. 4 E). The differences in T-ALL onset and incidence between the experimental groups in Figs. 1 and 4 correlated with the speed and level of T cell detection in the blood of the animals. _γ*c*_*^-/-^* hosts were faster at generating T cells, and this was most prominent in the group receiving pre-cultured HSPCs (Supplementary Fig.4). This suggested that culture conditions could affect HSPC function, but also that _γ*c*_*^-/-^* and *Rag2^-/-^*_γ*c*_*^-/-^* thymi differed in their immediate capacity to support T cell differentiation. To test the latter hypothesis, we injected 500 000 wild type HSPCs into non-irradiated _γ*c*_*^-/-^* or *Rag2^-/-^*_γ*c*_*^-/-^* hosts and analyzed them 3 or 9 weeks later (Fig. 4 F). Differences in total thymus cellularity were notorious between the two groups (Fig. 4 G). While thymopoiesis was evident in _γ*c*_*^-/-^* thymi by 3 weeks post injection, that was not the case in *Rag2^-/-^*_γ*c*_*^-/-^* thymi, which could not support thymopoiesis (Fig. 4 G, H). By 9 weeks post injection, both _γ*c*_*^-/-^* as well as *Rag2^-/-^*_γ*c*_*^-/-^* thymi showed CD4+CD8+ thymocytes, although total numbers were clearly lower in *Rag2^-/-^*_γ*c*_*^-/-^*hosts (Fig. 4 G). Thymopoiesis was accompanied by structural alterations that could be observed histologically (Fig. 4 I, J, K). While CD4+CD8+ double positive thymocytes occupied cortical areas, CD4+ or CD8+ single positive thymocytes located in the medullas (Fig. 4 I). The thymic epithelium is crucial for supporting thymopoiesis, and we could see obvious differences in epithelial composition and organization between _γ*c*_*^-/-^* and *Rag2^-/-^*_γ*c*_*^-/-^*(Fig. 4 I). While the first had clear cortical and medullary areas, this functional discrimination was absent in *Rag2^-/-^*_γ*c*_*^-/-^* mice and only emerged as the experiment progressed in time. No differences were detected in the level of bone marrow engraftment between the two host strains (Supplementary Fig. 5 A – D). Altogether, the data support that initial thymic epithelial structure determined the efficiency of thymopoiesis, and established T-ALL incidence.

**Figure 4.**
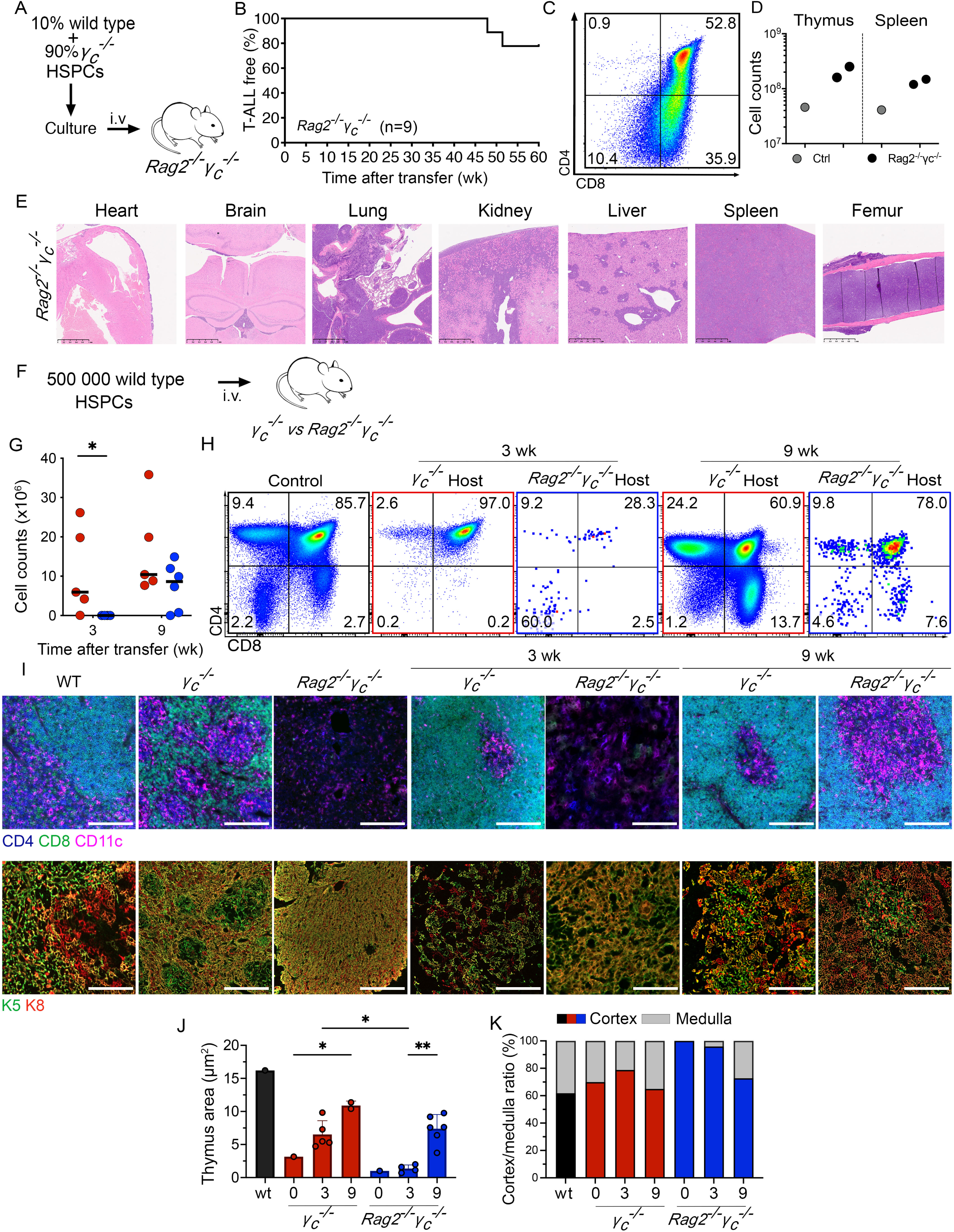
The thymic microenvironment determines T-ALL incidence and onset. A) 10^6^ HSPCs in a mixture of 10% wild type plus 90% _γ*c*_*^-/-^*, were cultured for 16 hours and i.v. injected into non-preconditioned *Rag2^-/-^*_γ*c*_*^-/-^* mice. CD45.1 and CD45.2 were used to distinguish donor/host-derived cells. B) Percentage of T-ALL free over time. C) Example of one T-ALL. Numbers show percentage of cells in each quadrant, gated on live donor cells. D) Cellularity of thymi and spleens, where each dot represents one mouse. E) Representative sections of the indicated organs stained with hematoxylin and eosin. Scale bars: 500μm. F) 500 000 wild type HSPCs were transplanted into non-preconditioned _γ_*_c_^-/-^* (n=10) or *Rag2^-/-^*_γ_*_c_^-/-^* (n=10) recipients and their thymi were analyzed 3 or 9 weeks after injection. One lobe was used for flow cytometry (H) and the other for histology (I). G) Thymus cellularity, where each dot represents one mouse, and the lines represent the medians. H) Representative examples of CD4CD8 profiles, pre-gated on donor cells. I) Immunohistology analysis of the indicated mice and timepoints. Pseudocolors are CD4 (blue), CD8 (green), CD4+CD8+ (cian), CD11c (magenta), keratin 5 (K5, green), keratin 8 (K8, red), K5+K8+ (yellow). Scale bar = 250 μm. J) Quantification of the thymus area (μm^2^) of the analyzed immunohistology sections. Each dot represents one thymus section, each bar shows the median. K) Percentage of cortex and medullary areas within each thymic section. Data in G-K are from 2 independent experiments containing n =4-6/experiment.

### Thymic microenvironment influences T-ALL development but shows limited impact on bulk transcriptional identity

Next, we performed bulk RNA-seq to characterize the transcriptomic landscape of T-ALL (Supplementary Fig. 6 A). Seven T-ALL samples were compared to wild-type thymocytes. Principal Component Analysis (PCA) of their gene expression showed clear separation between thymocyte control and leukemic cells, indicating that disease status is the primary driver of transcriptomic variation in the dataset (Supplementary Fig. 6 B, C). PCA did not resolve distinct subtypes among the leukemia samples (Supplementary Fig. 6 B). Consistently, unsupervised hierarchical clustering based on the 100 most variable genes in the T-ALL samples failed to identify major gene expression programs distinguishing the leukemias (Supplementary Fig. 6 D). Notably, several of the most variable genes were associated with sex (*e.g.*, *Eif2s3y, Ddx3y*, *Kdm5d*, *Xist*, *Tsix*). The prominence of sex-associated genes among the top sources of variability suggests that, beyond disease status, transcriptional differences between individual T-ALL samples are relatively subtle. Together, these results suggest a high degree of transcriptomic similarity between leukemias, irrespective of host or culture prior to i.v. injection. Nevertheless, as the number of samples is reduced, our interpretation is conservative. Differential gene expression analysis (DGEA) between T-ALL and control thymocytes identified significant upregulation of several membrane/erythrocyte-related genes, such as *Rhag*, *Car1*, *Add2* (Supplementary Fig. 6 E). We then specifically examined the expression of genes classically associated with T-ALL (*Lmo2*, *Lyl1*, *Tal1*, *Tfrc*, *Tlx1, Tlx3,* and *Hhex*). Among these, *Tal1* presented a strong upregulation, ranking among the top upregulated genes in T-ALL relative to controls (Supplementary Fig. 6 E, F). *Lmo2*, *Lyl1* and *Tfrc* were also significantly upregulated in T-ALL samples, although with more moderate effect sizes (Supplementary Fig. 6 F). In contrast, *Tlx1, Tlx3,* and *Hhex* exhibited very low expression levels across samples and were excluded during the low-expression filtering step of the analysis (see Methods, Supplementary Fig. 6 G), precluding reliable assessment of differential expression for these genes. Gene set enrichment analysis (GSEA)^35^ revealed significant enrichment in T-ALL of genes associated with cell proliferation, including E2F and Myc targets, and G2M checkpoint, as well as of metabolic gene sets such as Glycolysis, Oxidative Phosphorylation, and Fatty Acid Metabolism, all alterations commonly found throughout neoplastic transformation (Supplementary Fig. 6 H). In contrast, some gene sets related to immune signaling (*e.g.*, TNFα Signaling via NFκB and Allograft Rejection) were significantly depleted in T-ALL samples relative to control (Supplementary Fig. 6 H). Given the strong immune-related transcriptional programs characteristic of normal thymocytes, these changes reflected a relative reduction of immune signaling pathways in leukemic cells. Next, we compared the transcriptional profile of these T-ALL samples to samples obtained by thymus transplantation of wild type newborn thymus into *Rag2^-/-^*_γ_*_c_^-/-^* mice, as described previously^1, 10^ (Supplementary Fig. 6 I). Differences between thymus-graft-derived and those originating from limited bone marrow chimerism included higher expression of B-cell–associated genes in the latter, including immunoglobulin-related transcripts, consistent with the fact that adoptive transfer of wild type HSPCs also generated B cells, while thymus transplants generally fail to do so (Supplementary Fig. 6 J, K). Altogether, these data indicate that the thymic microenvironment plays a critical role in shaping T-ALL initiation and kinetics, with differences in epithelial organization between _γ_ *^-/-^* and *Rag2^-/--/-^* hosts profoundly influencing thymopoietic efficiency, leukemia onset, and disease incidence. Despite these contextual differences, transcriptomic profiling revealed that T-ALLs converge on a largely shared gene expression program that is clearly distinct from healthy thymocyte controls and dominated by canonical oncogenic, proliferative, and metabolic pathways, including strong activation of *Tal1*-associated transcription. Unsupervised analyses detected only limited heterogeneity among leukemias, suggesting that host environment and transplantation context modulate disease emergence and specific transcriptional features, without fundamentally altering the core leukemic program. Differences included the occasional presence of B-cell-associated signatures, resulting from the presence of B cells in the spleens of T-ALL resulting from limited bone marrow chimerism, where the samples were isolated from. Together, these findings support a model in which the initial thymic architecture dictates the efficiency and timing of leukemic transformation, while the resulting T-ALLs ultimately adopt a robust and conserved transcriptional state, highlighting both the importance of the niche during disease initiation and the need for higher-resolution approaches to uncover subtler layers of leukemia heterogeneity.

### Threshold for thymus autonomy inhibition

Thymus autonomy is the immediate consequence of impairing cell competition in the thymus^1^. Our data supports that limiting the frequency of bone marrow derived progenitors seeding the thymus enables thymus autonomy. As autonomy has a high risk of leukemia^10^, we sought to determine the threshold of bone marrow engraftment that establishes normal thymopoiesis and prevents thymus autonomy. We generated bone marrow chimeras using lethally irradiated _γ*c*_-deficient mice reconstituted with defined ratios of wild type to _γ*c*_*^-/-^* HSPCs (Fig. 5 A). In addition, we adoptively transferred 100% or 10% wild type plus 90% _γ*c*_*^-/-^* HSPCs (either cultured or not cultured) into non-irradiated _γ*c*_-deficient hosts (Fig. 5 A). Three weeks later, all mice were transplanted with wild type thymi and analyzed 5 weeks thereafter (Fig. 5 A). Taking CD4^+^CD8^+^ thymocytes as readout, we determined their origin and classified thymus grafts into three phenotypes: 1) exhausted, 2) in turnover, or 3) in thymus autonomy (Fig. 5 B). Exhausted thymi were those in which only _γ*c*_-deficient host cells (CD45.2^+^) were present. Thymus grafts in turnover were composed of wild type HSPC-derived thymocytes (CD45.1^+^), indicating that wild type bone marrow-derived progenitors replaced the thymus graft thymocytes. Finally, thymus autonomous grafts were those in which thymocytes of graft origin (CD45.1^+^/CD45.2^+^) persisted. Of note, autonomous thymopoiesis could be dominant in the thymus grafts or co-exist with bone marrow-derived thymopoiesis (Fig. 5 B). The level of bone marrow engraftment was consistent with the proportion of wild type HSPCs injected into lethally irradiated recipients, although some variation was observed (Fig. 5 C). In line with our previous observations, the level of bone marrow engraftment in non-irradiated recipients was extremely low (Fig. 5 C, zoomed in the dashed square). For both irradiated and non-irradiated mice, the frequency of thymus autonomy increased as the percentage of wild type engrafted HSPCs decreased (Fig. 5 D). While thymus autonomy was inhibited in irradiated hosts that received over 50% wild type HSPCs, in non-irradiated hosts thymus autonomy was prevalent even when all transplanted HSPCs were wild type (Fig. 5 D). The analysis of individual mice shows that even when the relative proportion of injected cells and the experimental conditions are the same, interindividual variability arises, especially when the percentage of wild type engraftment decreases (Supplementary Fig. 7 A), further supporting that there is stochasticity in thymus seeding. This is most noticeable among the non-irradiated individuals, where proficient thymus seeding progenitors are extremely reduced, and even _γ*c*_-deficient progenitors are capable of thymus seeding (Supplementary Fig. 7 A). Phenotypically, we previously defined thymus autonomy based on the percentage of thymus-graft CD4+CD8+ cells above 10%^32^ (Supplementary Fig. 7 B). Because the experiments here involved two sources of wild type thymocytes, i.e. thymus-graft and bone marrow-derived, we compared the distribution of samples with ≥ 10% thymus-graft CD4+CD8+ (grey circles) versus their representation within total CD4+CD8+ cells (Supplementary Fig. 7 C). We considered thymus autonomy those samples in which thymus-graft CD4+CD8+ cells were ≥ 10% of total CD4+CD8+ (Fig. 5 E, red circles). Importantly, such samples were among the ones with the highest absolute thymocyte counts of thymus-graft origin, suggesting a higher risk for leukemic transformation (Fig. 5 F). Taken together, our data supports that the relative proportion of wild type HSPCs injected to correct _γ*c*_-immunodeficiency should be homogeneous to reduce the risk of chimerism and favor the proportion of _γ*c*_-proficient progenitors engrafted. Further, conditioning the host prior to HSPC injection optimizes progenitor engraftment, thereby reducing thymus autonomy.

**Figure 5.**
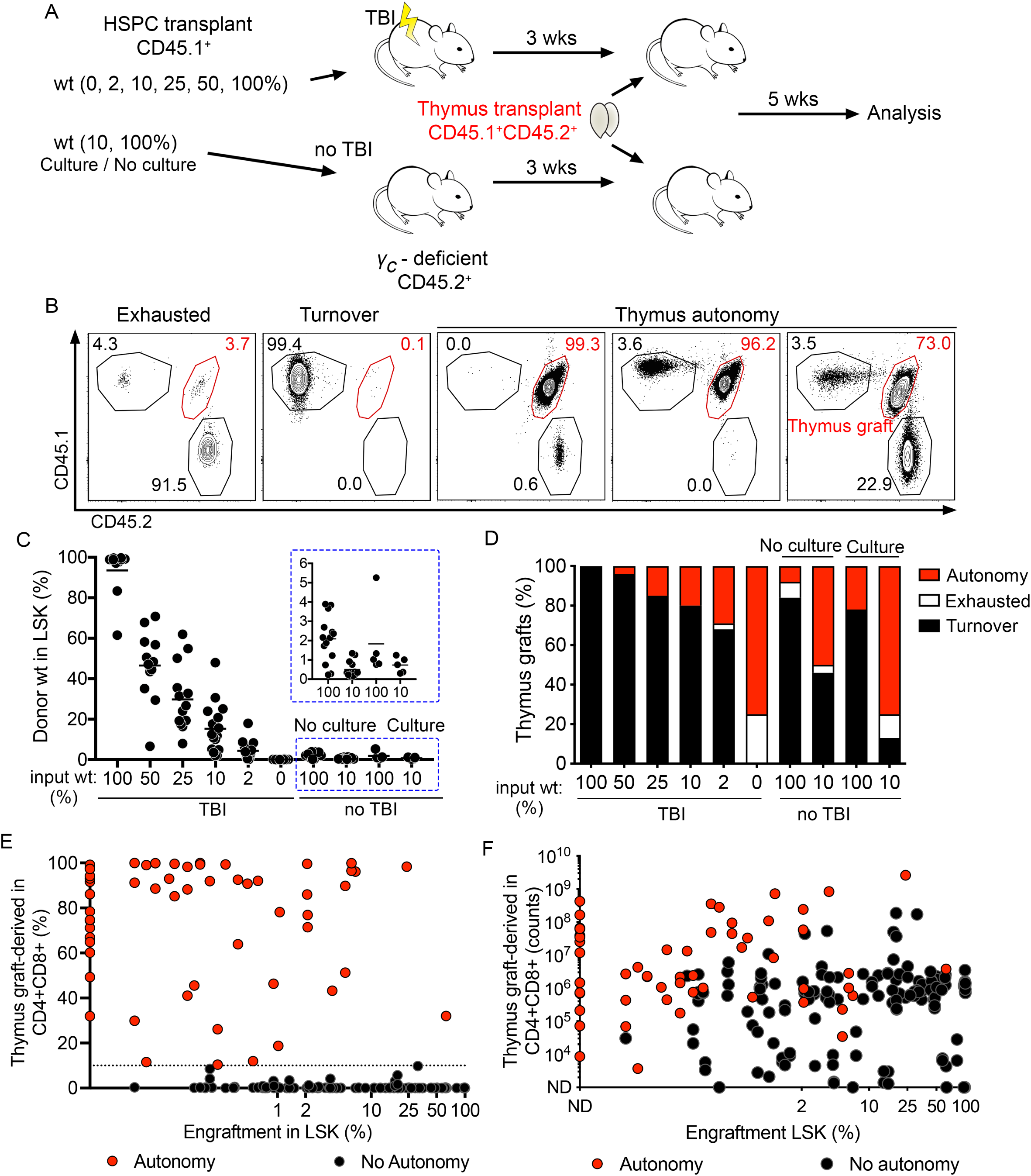
Threshold for inhibition of thymus autonomy. A) 10^6^ HSPCs in the indicated percentages of wild type (CD45.1^+^) to _γ*c*_*^-/-^* (CD45.2^+^) cells were i.v. injected into hosts (CD45.2^+^) that were either previously lethally irradiated (total body irradiation, TBI) or untreated. For the non-irradiated hosts, one group received pre-cultured cells, and the other non-cultured cells. Three weeks later, mice were transplanted with newborn wild type thymi (CD45.1^+^/CD45.2^+^) and analyzed 5 weeks thereafter. B) Representative examples of phenotypic outcomes depicting the origin of cells. C) Effective level of engraftment in bone marrow progenitors measured in LSK. Blue highlighted region in the main graph is zoomed in above. D) Percentage of thymi exhausted, autonomous or in turnover in the indicated groups. E) Relation between effective engraftment in bone marrow LSK (x-axis) and the percentage of thymus graft-derived CD4CD8 double positive thymocytes. F) Relation between absolute cell counts of thymus graft-derived CD4CD8 double positive thymocytes and engraftment in bone marrow LSK. Shown is cumulative data from 5 independent experiments where hosts were _γ*c*_*^-/-^* (n=44) or *Rag2^-/-^*_γ*c*_*^-/-^* (n=71) in total.

## DISCUSSION

Here we show that inefficient _γ*c*_-deficiency correction resulting in low level of wild type bone marrow chimerism generates discrete waves of thymopoiesis with intermittent T cell production and export. This was detected in the blood by measuring recent thymic emigrants, which showed that thymus activity occurred in confined time periods that were interspersed by intervals without T cell production. Such observation can only be explained by sporadic events of thymus colonization by wild type hematopoietic progenitors. The estimates for thymus seeding progenitors in wild type mice are in the range of 10 cells per day^36^, and each progenitor that reaches the thymus creates one wave of thymopoiesis. The time between two consecutive seeding events is predictably short and ensures constant thymopoiesis. In _γ*c*_-deficient mice with a low-level reconstituted bone marrow, the number of proficient thymus seeding progenitors is expectably lower. Thus, the time between two consecutive waves of wild type thymopoiesis will vary due to stochasticity in thymus seeding, and this impairs thymic cell competition and enables DN3-early self-renewal. Thymus autonomy is thereby triggered and can possibly only be suppressed if another wave of thymopoiesis supersedes the thymus before transformation occurs.

In such a scenario, thymus analyses provide snapshots of thymus waves. We consistently detected phenotypic variability documenting waves of thymopoiesis at different stages, as well as thymi with only circulating T cells. Of note, the most frequent phenotype within the CD4-CD8-double negative (DN) stages lacked the most immature ETP to DN2 populations and contained DN3 thymocytes, in line with the capacity of the latter to self-renew^32^. The competition of DN3 thymocytes for thymic niches, and their dependency from interleukin 7 receptor (IL-7r) is well-known in *IL-7r*_α_*^-/-^* mice, which have open niches that can be directly colonized by progenitors injected into the blood stream without the need of irradiation^37^. We previously showed that IL-7r regulates cell competition non-cell autonomously^1, 7^. Further, blocking thymus seeding by thymopoiesis-proficient progenitors hinders cell competition, and triggers thymus autonomy ^8, 9^, which later causes T-ALL^1, 10, 30^. This is in line with our current data showing that if thymus seeding is sporadic, the number of thymocytes generated is too low to outcompete older DN3 thymocytes with a longer time of thymus residency. In parallel, aberrant CD4+CD8+ thymocytes lacking expression of TCRβ emerge prior to T-ALL onset. Further studies will address how that population expands to generate T-ALL.

Our data is relevant to understand possible consequences of correcting immunodeficiency of SCID-X1 patients^38^. The current standard of care of these patients is a transplant of healthy bone marrow, and no adverse effects have been reported^19, 21, 39^. In the absence of compatible donors, gene therapy might be a solution, as is for another type of SCID, the ADA-SCID^40, 41^. However, the absolute number, as well as the relative proportion, of gene therapy-corrected cells transplanted is critical. The efficiency of gene correction of HSPCs will influence the level of chimerism in the host. This means that there is a risk of infusing back a heterogeneous population of progenitors that can establish a suboptimal level of chimerism, instead of fully correcting immunodeficiency. Importantly, gene therapy requires induction of cell proliferation to transduce HSPCs. Although the goal is to expand cell number without losing multipotency, cell culture conditions are likely to interfere with HSPC function. In line with this possibility, T cells were produced faster in a SCID-X1 patient cohort treated by gene therapy as compared with a cohort receiving haploidentical hematopoietic stem cell transplantation^42^. Our data show that mouse HSPCs cultured overnight change, accelerating T-ALL onset and increasing incidence. This suggests that cultured HSPCs are less effective to home to the bone marrow, that they lose some multipotency and/or generate more differentiated progeny. In either case, if such changes occur in a gene therapy setting, then the effective number of HSPC with capacity for long term engraftment is probably reduced. Finally, although opening the bone marrow niches bears significant risks for patients, it also ensures more efficient engraftment, and therefore lower risk of thymus autonomy and leukemia.

It is noteworthy that the thymocyte population involved in cell competition is the same that has self-renewal potential. Such property is likely to have a physiological function that we have not yet found. One possibility could be that DN3-early self-renewal potential is associated to β-selection, optimizing the probability to generate a productive TCRβ rearrangement. In any case, DN3-early cells have been reported to self-renew prior to full T-ALL onset in transgenic mouse models induced by either *Lmo2*^43^, or *Lmo1* plus *Tal1*^44^ overexpression. This property seems to confer the DN3-early population with characteristics of a super-competitor^45^, suggestive that thymocyte self-renewal could be a common mechanism underlying T-ALL initiation^46^. In fact, *LMO2* activation is considered an initiating event in human T-ALL^47^. These examples point at the DN3-early as a population that must be kept in check, and we consider that cell competition is crucial in that respect. At steady state, young DN3-early are constantly outcompeted by young cells with less time of thymus residency. Those with shorter time of thymus residency are more fit than the ones dwelling in the thymus for longer time. If, however, these young precursors do not reach the thymus in sufficient numbers to outcompete the older DN3-early, the latter persist, self-renew, and cause leukemia. Further work will require uncovering the molecular regulation of cell competition in the thymus. Interestingly, thymus autonomy has been reported previously following intrathymic injection of bone marrow progenitors into *Zap70^-/-^*mice without pre-conditioning^48, 49^. These mice maintained thymopoiesis autonomously for 28 weeks without ever developing malignancy. Opposite to all our observations of thymus autonomy^1, 8, 10, 32^, donor HSPCs injected in *Zap70^-/-^*generated all thymocyte populations, including the early thymus progenitor (ETP)^48, 49^. This suggests that in those experiments cell competition at the DN3 stages is probably maintained, thereby preventing malignant transformation. Furthermore, the niches occupied by the injected cells must differ significantly between the different experimental settings.

Collectively, our data reveal that DN3-early thymocytes possess an intrinsic capacity for self-renewal and highlight the physiological cost of this property when bone marrow correction of _γ*c*_-deficiency falls below the threshold required to restore effective thymopoiesis. It should be noted that adult hematopoietic progenitors are capable to generate thymocytes with self-renewal potential, as our former work had only demonstrated this property in newborn thymocytes. Using a mouse model of limited bone marrow reconstitution, we show that cell competition in the thymus is a fundamental quality control mechanism that suppresses malignant transformation. Ultimately, our findings indicate that any condition impairing thymic cell competition and triggering thymus autonomy has a high risk of malignant transformation and leukemia.

## ONLINE METHODS

### Mice

All mice were maintained and bred at Gulbenkian Institute for Molecular Medicine (Oeiras), former Instituto Gulbenkian de Ciência. C57BL/6J (CD45.2^+^) were in a colony frequently refreshed by mice imported from Charles River. B6.SJL-Ptprc3^a^ (CD45.1^+^, here termed B6.SJL), stock #002014, were purchased from The Jackson Laboratory and kept in a colony at IGC. _γ*c*_*^-/-^* ^31^, JAX stock #003174 were purchased from The Jackson Laboratory. *Rag2^-/-^*_γ*c*_*^-/-^*mice were obtained by crossing *Rag2^-/-^*^50^ and _γ*c*_*^-/-^* mice, both in B6 background, as previously described^10^. Mice of both genders were used at an age of 4 to 8 weeks as recipients and kept in individually ventilated cages in specific pathogen-free (SPF) conditions. All experiments were conducted in compliance with Portuguese and European laws and approved by the internal Animal Welfare Office (ORBEA-GIMM) and *Direção Geral de Alimentação e Veterinária (DGAV)*.

### Hematopoietic stem and progenitor cell transplantation

Bone marrow cells from 6- to 8-week-old B6.SJL or B6 (CD45.1^+^ or CD45.2^+^, wild type) or *Rag2^-/-^*and _γ*c*_*^-/-^* (CD45.2^+^ or CD45.1^+^) were incubated with biotinylated antibodies against CD3, CD4, CD8, CD11b, CD11c, CD19, Gr-1, NK1.1 and Terr119 and depleted of lineage*^+^* cells by magnetic separation using Biotin Binder Dynabeads (Thermo Fisher Scientific). Depending on the experiment, cells were either directly injected intravenously, or pre-cultured and then injected into the tail vein. Where indicated, cells were cultured for 16 hours at one million lineage-depleted cells per well in 24-well plates in StemPan Serum-Free Expansion Medium (Stem cell technologies, 096000) containing 200 ng/mL vaccinia virus B18R protein (Biovision, 8016-10 for the first set of experiments, AcroBiosystems, B1R-H52H6 for the latest, without noted differences) and the following cytokines from PeproTech: Flt3L (100 ng/mL), SCF (100 ng/mL), TPO (50 ng/mL) and IL-3 (20 ng/mL). Injected cells consisted of a mixture of a total of 1×10^6^ cells composed of 10% wild type (CD45.1^+^ or CD45.2^+^) and 90% _γ*c*_*^-/-^* (CD45.1^+^ or CD45.2^+^) cells. Recipient mice were _γ*c*_*^-/-^* (CD45.1^+^ or CD45.2^+^) or *Rag2^-/-^*_γ*c*_*^-/-^* (CD45.1^+^ or CD45.2^+^). Mice were used as CD45.1^+^ or CD45.2^+^ for donor/host discrimination, without noted functional differences for whether donors were CD45.1^+^ or CD45.2^+^. For Fig. 3, Femurs and tibias from Tg(*Rag-GFP)* mice^33^ were shipped from ICVS, Braga, to IGC at 4°C overnight and bone marrow single cell suspension were prepared and transplanted in the morning for the 1^st^ experiment. For experiments depicted in Fig. 5, mice were preconditioned by gamma-irradiation (600 rad), as indicated, prior to injection.

### Blood analyses

Mice were periodically bled from the submandibular vein to assess peripheral reconstitution and/or to screen for leukemia. Blood was collected into PBS with 1.6 mg/ml EDTA and erythrocytes were lysed using Gey’s solution (130.9 mM NH□Cl, 5.0 mM KCl, 0.84 mM Na□HPO□, 0.18 mM K□PO□·2H□O, 5.55 mM Glucose, 0.028 mM phenol red, 1.03 mM MgCl□·6H□O, 0.28 mM MgSO□·7H□O, 1.53 mM CaCl□·2H□O, 13.4 mM NaHCO□ before staining with appropriate antibodies.

### Thymus transplants

Thymi from newborn mice were collected and thymic lobes were split. Recipient mice were grafted under the kidney capsule with two lobes, and placed into opposing extremities of the kidney. Mice were kept anesthetized with Xylazine (16 mg/kg) and Ketamine (100 mg/kg). *Rag2^-/-^*_γ*c*_*^-/-^* or _γ*c*_*^-/-^* (CD45.2^+^) recipients of both genders with 5 to 7 weeks were used and thymus donors were wild type (CD45.1^+^CD45.2^+^).

### Flow cytometry

Organs were collected and cell suspensions were prepared in PBS with 2% FBS. Cells were blocked with mouse IgG (Jackson laboratories) and then stained with the antibodies against the following: CD3 (clone 145-2C11; conjugated to APC-Cy7), CD4 (GK1.1; PE, PE-Cy7, BV421, BV605 or BV650), CD8 (53-6.7; FITC, PerCP-Cy5.5, APC Fire 750 or BV711,), CD11b (M1/70; APC), CD19 (6D5; FITC or PE-Cy7), CD25 (PC61; BV605 or FITC), CD44 (IM7; BV711, eFluor450, FITC or PerCP-Cy5.5), CD48 (HM48-1; APC). CD71 (RI7217; FITC), CD150 (TC15-12F12.2; BV605), Gr-1 (RB6-8C5; PE or APC), Kit (2B8; APC or APC-Cy7), Ly6D (49-H4; FITC), Flt3 (A2F10; PE), IL-7r (A7R34; PE-Cy7), Sca-1 (D7; PercP-Cy5.5), CD45.1 (A20; FITC, PE, PE-Cy7, APC, BV421 or BV650) and CD45.2 (104; FITC, PE, APC, APC-Cy7, Pacific Blue or BV711), all from BioLegend. The lineage mix consisted of CD3 (145-2C11), CD4 (GK1.1), CD8 (53-6.7), CD11b (M1/70), CD11c (N418), CD19 (6D5), Gr-1 (RB6-8C5), NK1.1 (PK136) and Ter119 (TER-119), all conjugated to biotin and from BioLegend. Biotinylated antibodies were recognized by streptavidin conjugated to BV785 (BioLegend). Dead cell exclusion was performed using Zombie Pacific Orange (BioLegend). Intracellular staining against Ki-67 (16A8; AF700) and TCRβ (H57-597; AF647 or BV711), from BioLegend was performed with True-Nuclear Transcription Factor Buffer Set (BioLegend). Samples were acquired in a BD LSRFortessa X-20 analyzer and analyzed on FlowJo 10.10.0.

### Histopathology

Histological analysis was performed at the Histopathology Facility of the GIMM Institute. Briefly, samples were fixed in 10% neutral-buffered formalin for a minimum of 48 hours at room temperature, except the brain that was fixed for 1 week. Samples were processed automatically (Leica HistoCore Pearl), embedded in paraffin, and sectioned at 3 µm. Sections were mounted on glass slides and stained with hematoxylin and eosin (H&E) following standard protocols. Slides were analyzed with a DMLB2 microscope (Leica), and images were acquired with a DFC320 camera (Leica) and NanoZoomer-SQ Digital slide scanner (Hamamatsu Photonics).

### Immunohistology

Thymi were collected and cryopreserved in OCT. Sections (8 µm thick) were obtained using a Leica Cryostat CM 3050 S, dehydrated with acetone and stored at -80 °C until use. Sections were rehydrated, incubated with DAPI and mouse IgG, stained overnight with rabbit Keratin 5 (Poly19055), Keratin 8-Alexa647 (1E8), CD11c-Bio (N418), CD4-Alexa647 (GK1.5) and CD8-Alexa488 (53-6.7), followed by Streptavidin-Cy3 and anti-rabbit Alexa488, as appropriate. Slides were mounted with Fluoromount G medium (Invitrogen) and images were acquired on a Zeiss Axio Imager with Apotome 2 microscope. Fiji 2.14.0/1.54f was used for image treatment and QuPath-0.6.0-x64 for analysis and quantification.

### Statistical Analyses

We performed Cox proportional hazards regression with a time-dependent coefficient to evaluate whether the effect of culture condition on the hazard varies over time. The model was specified as:

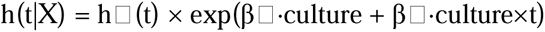

where h(t|X) is the hazard function at time t, h□(t) is the baseline hazard function, culture is a binary indicator variable, and t represents time (in weeks). The interaction term (culture×t) allows the log-hazard ratio for culture to change linearly with time, testing whether the culture effect strengthens or weakens over the follow-up period. The model was fitted using the lifelines CoxPHFitter implementation (v0.27+) with duration measured in weeks as the time-to-event outcome and a binary event indicator. Statistical analysis from figures 2 and 4 were performed with GraphPad, Prism 10.6.1.

### RNA isolation and sequencing

Cell pellets containing 5 million or 20 million cells were prepared from leukemia samples and stored as dry pellets at −80°C until use. Total RNA was then isolated by lysing the cells with Trizol (Invitrogen), followed by RNA phase separation with trichloromethane/chloroform (Carl Roth), precipitation with 2-propanol (VWR BDH Chemicals), and dilution in RNAse-free water. Quality control and RNA concentration was determined in a Fragment analyzer. The isolated RNA were stored at −80°C until sequencing. Sequencing was performed by Novogene Co., Ltd.. For sequencing, first messenger RNA was purified from total RNA using poly-T oligo-attached magnetic beads. After fragmentation, the first strand cDNA was synthesized using random hexamer primers followed by the second strand cDNA synthesis. The library was ready after end repair, A-tailing, adapter ligation, size selection, amplification, and purification. The library constructed was checked with Qubit and real-time PCR for quantification and bioanalyzer for size distribution detection. Quantified libraries were pooled, based on the effective concentration and targeted data amount, and sequenced on Illumina platforms.

### RNA-seq analyses

Quality control of raw sequencing reads (in FASTQ format) was performed using multiQC (v1.17)^51^ with default parameters. Raw reads were then trimmed using Trimmomatic (v0.39)^52^ operating in paired-end (PE) mode, with appropriate adapter sequences and parameters. Trimmed reads were aligned to the mm39 reference genome using STAR (v2.7.9a)^53^ with the following parameters:

- - outFilterType BySJout \
- -alignSJoverhangMin 8 \
- -alignSJDBoverhangMin 1\
- -alignIntronMin 20 \
- -alignIntronMax 1000000 \
- -alignMatesGapMax 1000000 \
- -outFilterMultimapNmax 1 \
- -outFilterMismatchNmax 999 \
- -outFilterMismatchNoverLmax 0.04 \

Aligned reads (BAM files) were subsequently annotated using the GENCODE vM38 primary assembly^54^ and quantified with featureCounts (v2.0.3)^55^.

All the following analyses were performed in R (v4.1.2). For each gene, raw read counts were first summed across samples, with genes with total log_10_ read counts < 3.5 removed, to filter out low-expression genes (Supplementary Fig. 4G). Filtered read counts were then used to calculate library size scaling factors using the trimmed mean of M-values (TMM; calcNormFactors function from edgeR (v3.36.0))^56, 57^. These were subsequently normalized and log_2_-transformed using the voom function from limma (v.3.50.3)^58, 59^.

For the DGEA between T-ALL and WT, we calculated the log2 FC of the means. Standard errors of the means were then moderated using empirical Bayes shrinkage^60^. The log_2_ FC of the means and the Bayesian log-odds of differential expression (the so-called B-statistic) □Ref 10] were the chosen statistics for differential gene expression, as indicated on the volcano plot in Supplementary Fig. 4C.

For hierarchical clustering, the 100 genes with the most variable expression across T-ALL samples were selected. Using pheatmap package (v1.0.13)^61^, pairwise Euclidean distances were computed between samples based on the expression of the selected genes, and clusters were derived using the Ward.D2 linkage method^61^. The resulting dendrogram was visualized together with a heatmap (Supplementary Fig. 4D).

GSEA was performed using the fgsea (v1.20.0) R package^35^. Genes were ranked by t-statistics from DGEA, ordered from most positive to most negative, and analyzed against the Hallmark Pathways gene sets^62^. The normalized enrichment score (NES) was extracted to quantify the extent to which a given gene set is enriched at the extremes of the ranked gene list ^35, 62^. A positive NES indicates that the gene set is enriched amongst over-expressed genes, whereas a negative NES indicates its depletion (*i.e.*, its enrichment amongst under-expressed genes).

## ACKNOWLEDGEMENTS

This work was supported by the “La Caixa” Foundation grant LCF/PR/HR22/52420023 to VCM, the Portuguese National Research Council (Fundação para a Ciência e Tecnologia [FCT] Grant PTDC/MED-IMU/3649/2021 to VCM and Instituto Gulbenkian de Ciência (IGC) of the Calouste Gulbenkian Foundation. VCM was supported by an individual contract awarded by FCT (CEECIND/03106/2018). CCP is a PhD student of the IGC Integrative Biology and Biomedicine (IBB) PhD Program and is supported by an individual FCT PhD Fellowship ref. UI/BD/154901/2023. RAP and CVR were PhD students of the IBB PhD Program and were supported by individual FCT PhD Fellowships refs. PD/BD/114341/2016 and PD/BD/139190/2018, respectively. FXB and AK are PhD Students of the CAML PhD Program and supported by individual FCT PhD Fellowships refs. UI/BD/154080/2022 and UI/BD/153363/2022, respectively. We thank Jonathan Howard and Nuno L. Barbosa-Morais for critical reading and contributing for the final version of the manuscript. We thank Margarida Correia-Neves for providing the *Tg(Rag2p-GFP)* bone marrow cells. We thank Olivier Joffre and Pamela Fink for the *Tg(Rag2p-GFP)* mice. We thank the team of the Animal House, Flow Cytometry, Microscopy, Histopathology, and Genomics and Advanced Data Analysis Units of GIMM in supporting this work.

## AUTHOR CONTRIBUTIONS

BSPO and RAP designed and performed experiments, analyzed data and wrote the manuscript. CCP performed experiments and analyzed RNA seq data; RVP, CVR, SA performed and analyzed experiments; MXGP performed RNA experiments. FXB, AK compared transcriptomes between T-ALL subgroups. P.F. did the histopathological assessment of the leukemic samples, T.P. did the preliminary bioinformatics analysis of the RNA-seq data. VCM conceived the study, designed research and experiments, analyzed data and wrote the manuscript. All authors edited and contributed to the final version of the manuscript.

