## Supplementary Figures for "Limited bone marrow chimerism impairs cell competition in the thymus and causes leukemia"

Supplementary Figure 1

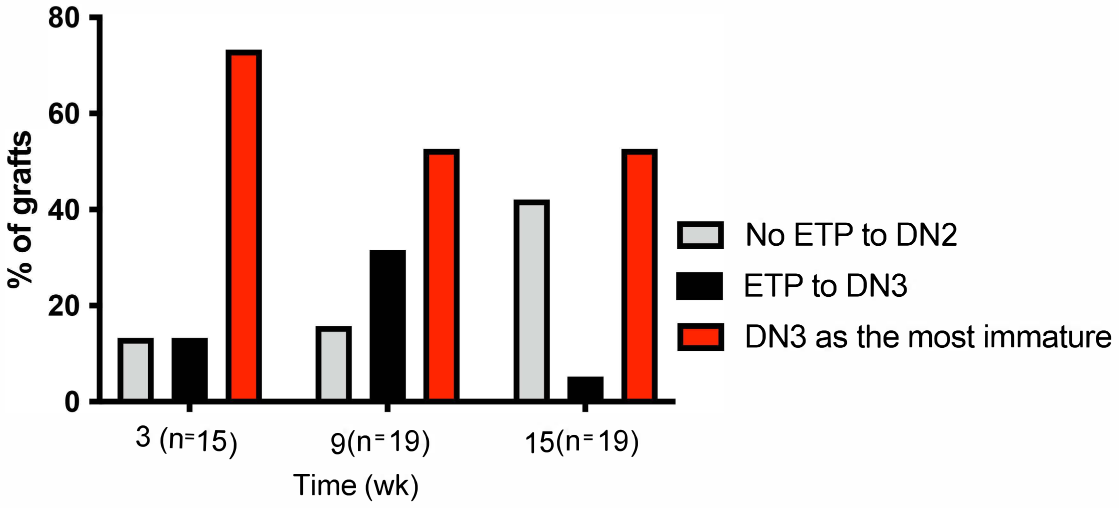

### Supplementary Figure 2

A

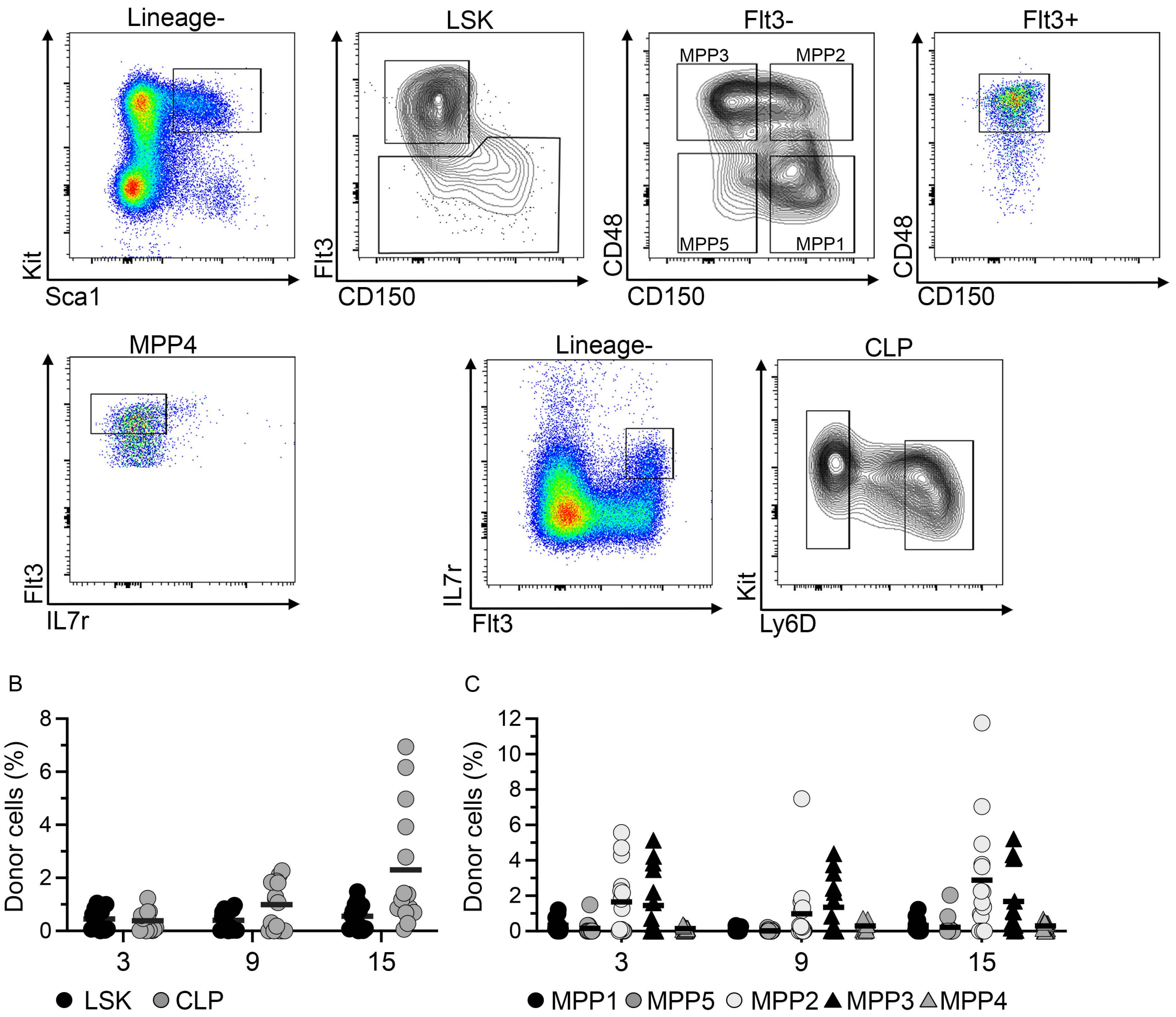


D

| Time (wk) | LSK | CLP | MPP1 | MPP5 | MPP2 | MPP3 | MPP4 |
|---|---|---|---|---|---|---|---|
| 3 | 0.45 ± 0.41 | 0.38 ± 0.37 | 0.33 ± 0.38 | 0.16 ± 0.39 | 1.66 ± 1.92 | 1.45 ± 1.85 | 0.15 ± 0.11 |
| 9 | 0.40 ± 0.37 | 0.99 ± 0.89 | 0.11 ± 0.11 | 0.02 ± 0.06 | 0.98 ± 1.85 | 1.35 ± 1.53 | 0.29 ± 0.24 |
| 15 | 0.56 ± 0.49 | 2.30 ± 2.28 | 0.02 ± 0.06 | 0.23 ± 0.57 | 2.88 ± 3.27 | 1.67 ± 1.90 | 0.29 ± 0.23 |

### Supplementary Figure 3

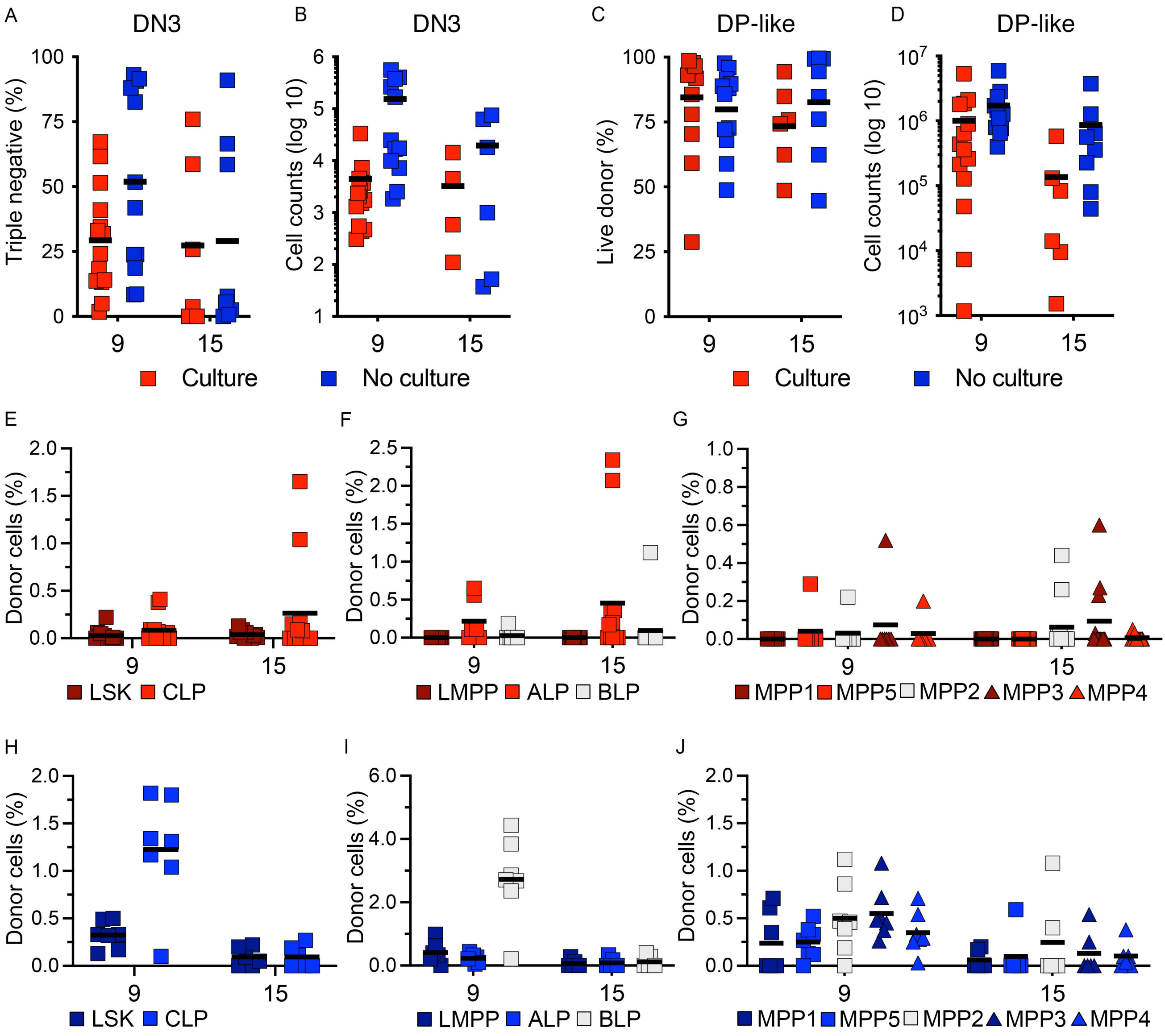

Supplementary Figure 4

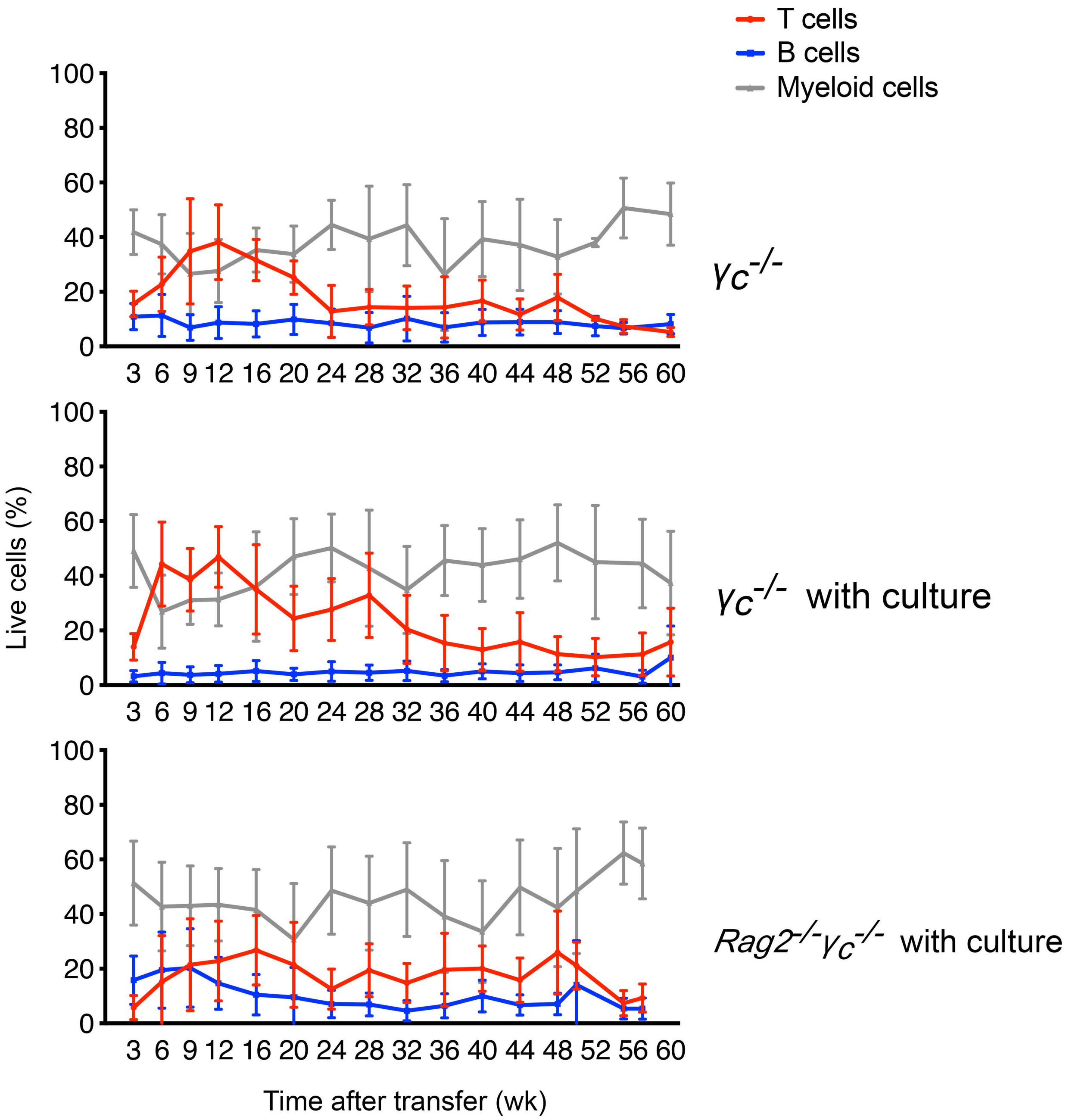

### Supplementary Figure 5

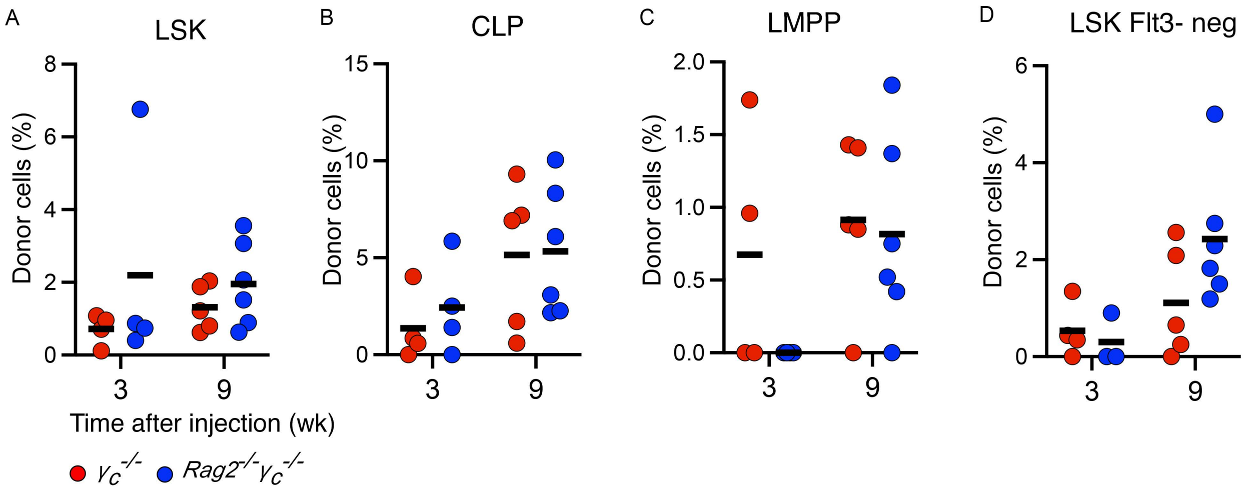

### Supplementary Figure 6

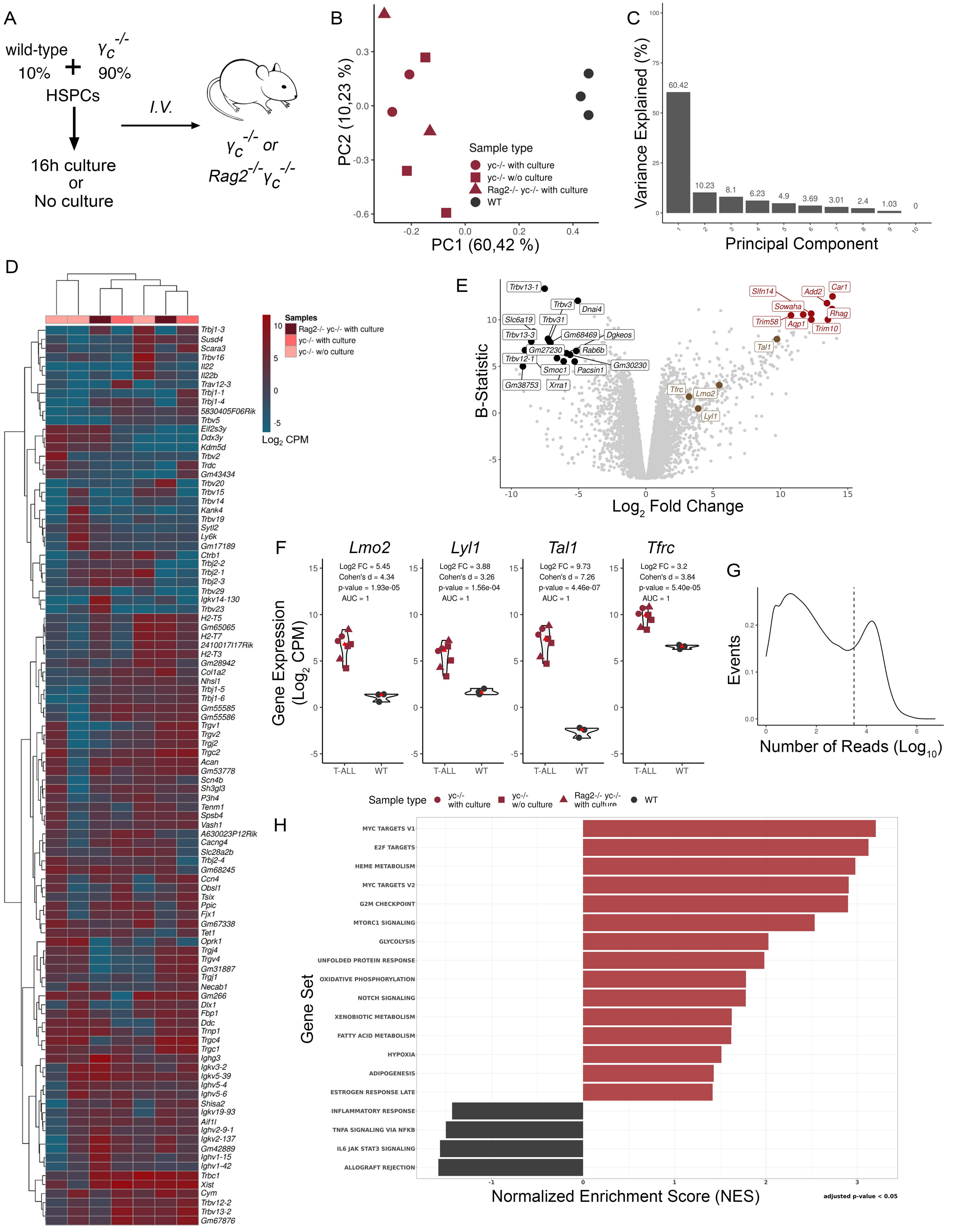

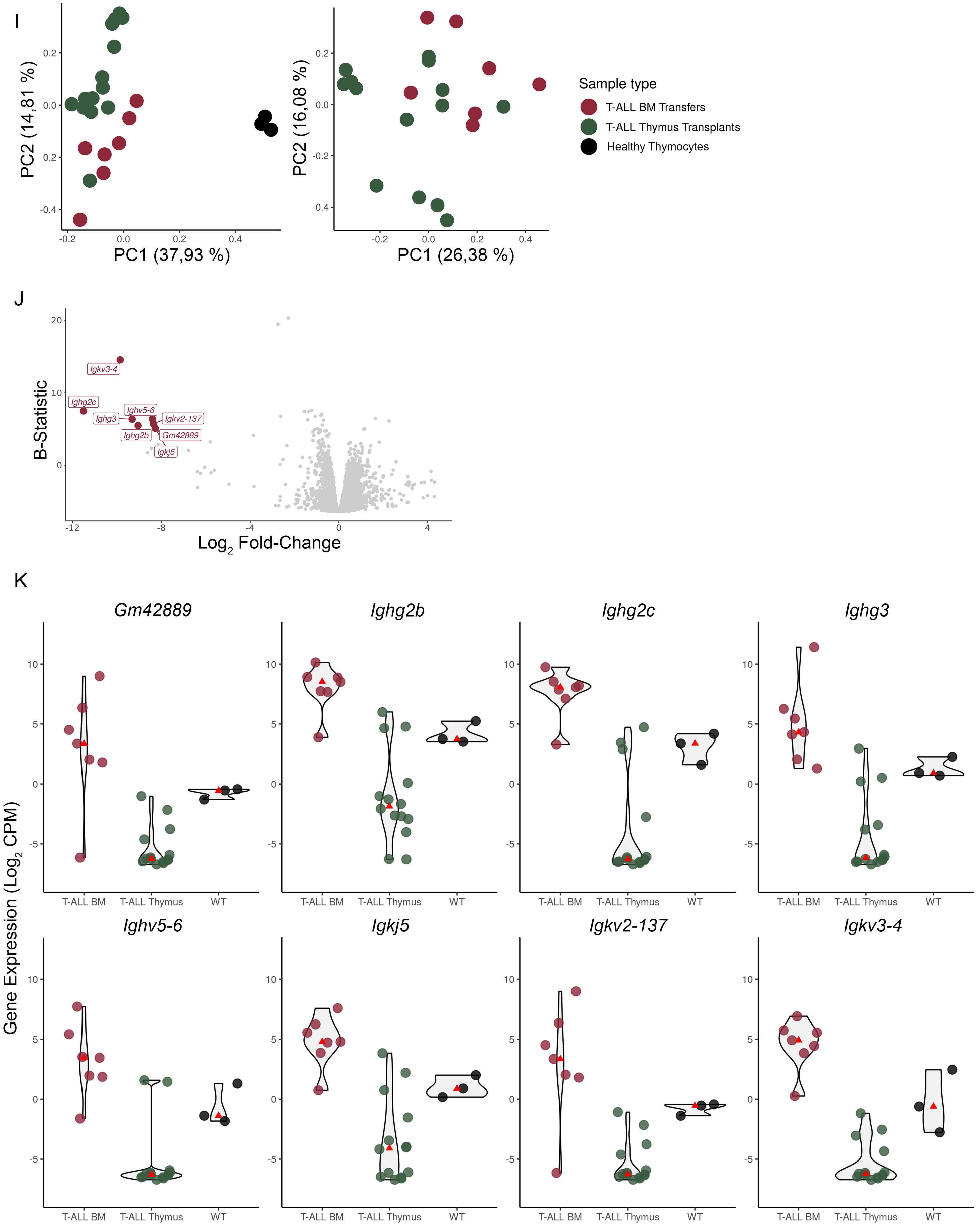
I
PC1 (37,93 %)
PC2 (14,81 %)
PC1 (26,38 %)
PC2 (16,08 %)
Sample type
T-ALL BM Transfers
T-ALL Thymus Transplants
Healthy Thymocytes
J
B-Statistic
Log2 Fold-Change
Igkv3-4
Ighg2c
Ighv5-6
Ighg3
Igkv2-137
Ighg2b
Gm42889
Igkj5
K
Gene Expression (Log2 CPM)
Gm42889
Ighg2b
Ighg2c
Ighg3
Ighv5-6
Igkj5
Igkv2-137
Igkv3-4
T-ALL BM
T-ALL Thymus
WT

Supplementary Figure 7

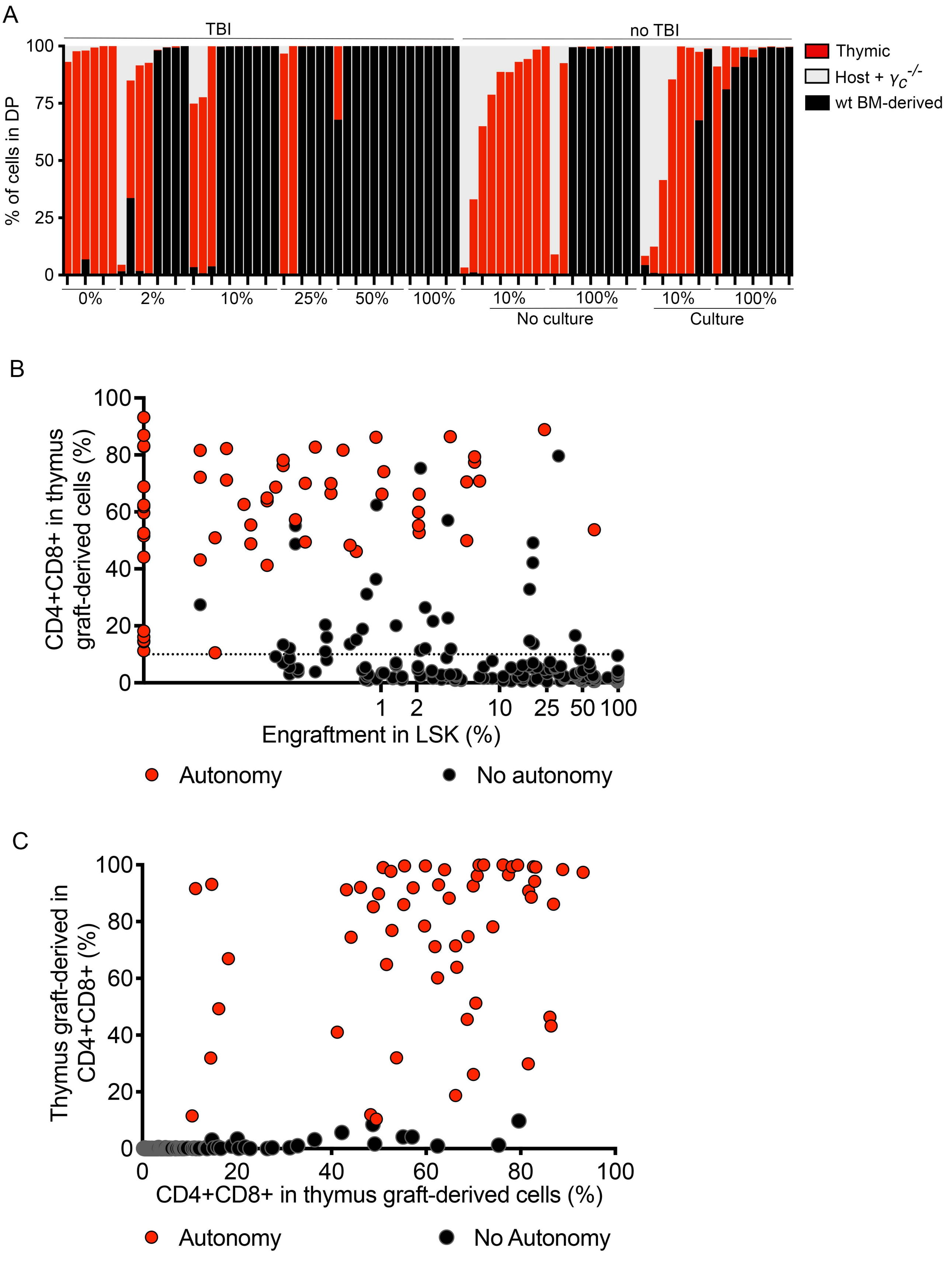

#### SUPPLEMENTARY FIGURE LEGENDS

**Supplementary Figure 1. Quantification of phenotypes within DN thymocytes.** Thymi analyzed in Fig. 2. D. were quantified for the presence of donor ETP, DN2, and DN3 as shown. Only thymi with wild type donor CD4/CD8 double positive thymocytes, corresponding to thymi represented by the black bars in Fig. 2. B. were considered in this analysis.

**Supplementary Figure 2. Quantification of donor-derived cells in bone marrow progenitors.** The bone marrow progenitors of mice in Fig. 2 were analyzed for the presence of cells of wild type (injected) origin. A) Gating strategy. B, C) Quantification of donor-derived cells in the indicated populations. Each dot represents a mouse and the line represents the mean. D) Mean ± Standard Deviation of the data shown in B, D.

**Supplementary Figure 3. Quantification of thymus and bone marrow cells.** Mice in Fig. 3 A were analyzed for the A) percentage and B) cell counts of DN3, and C) percentage and D) cell counts of CD4+CD8+ double positive-like (DP-like) cells. (E to J) The percentage of the indicated donor populations in live bone marrow cells was determined for the mice that received cultured HSPCs (E-G, plots with red datapoints) and for the mice that received non-cultured HSPCs (H-J, plots with red datapoints). Each square represents a mouse and the line is the mean.

**Supplementary Figure 4. Blood analysis**. Time course of blood analysis of the mice shown in Fig. 1 and Fig. 4A-E. Cells analyzed were live CD3+ (T cells), CD19+ (B cells), or Gr1+CD11b+ (Myeloid).

**Supplementary Figure 5. Quantification of bone marrow engraftment**. Mice in Fig. 4 F were analyzed for the percentage of donor cells in A) LSK, B) CLP, C) LMPP, and D LSK Flt3-negative. Red dots represent data from $\gamma_C^{-/-}$ hosts, and blue dots from *Rag2*$^{-/-}$ $\gamma c^{-/-}$ hosts.

**Supplementary Figure 6. Characterization of T-ALL transcriptomes. A)** Experimental design. **B)** PCA of the normalized gene expression in T-ALL samples (red, n = 7) and WT thymocytes (black, n = 3). Symbols and colors reflect the origin of the samples. **C)** Scree plot showing percentage of variance explained by each principal component. **D)** Heatmap and dendrogram of gene expression ($\log_2$ CPM) across T-ALL samples. Columns correspond to samples and rows to the 100 most variable genes across T-ALL samples. **E)** Volcano plot of DGE between T-ALL and WT thymocytes (X-axis – $\log_2$ fold change (FC); Y-axis – B-statistic). Genes are shown as circles. Most differentially expressed transcripts are colored red if upregulated (arbitrarily, $\log_2$ FC > 10 and B-statistic > 10) and black if downregulated (arbitrarily, $\log_2$ FC < -5 and B-statistic > 5). T-ALL-related genes are colored in brown. **F)** Expression of selected T-ALL-related genes (Y-axis – $\log_2$ CPM) in T-ALL samples (dark red) and WT thymocytes (black). Symbols and colors reflect the origin of the sample. The bright red triangle represents median expression. $\text{Log}_2$ FC from each gene's differential expression analysis, Cohen's d (as an effect size metric), two-sided t-test p-value and area under the ROC curve (AUC) are shown on the plot. **G)** Density plot of the distribution of the per-gene sum of RNA-seq reads across samples, illustrating the low-expression gene filtration step. The dashed vertical line (x = 3.5) represents the minimum number of total reads (3162) across samples necessary for a gene to be kept for the subsequent analyses. **H)** GSEA of Hallmark gene sets in DGE between T-ALL and WT thymocytes. Only gene sets with an adjusted p-value < 0.05 are shown. **I)** PCAs of the normalized gene expression in T-ALL samples resulting from thymus transplants (red, n = 14), BM transfers (green, n = 8) and (left) WT thymocytes (black, n = 3). **J)** Volcano plot of DGE between T-ALLs originated from thymic transplantation and BM transfers (X-axis – $\log_2$ FC; Y-axis – B-statistic). Genes are shown as circles. Most differentially expressed transcripts are colored dark red if downregulated (arbitrarily, $\log_2$ FC < -6 and B-statistic > 5). **K)** Expression of selected differentially expressed genes, as identified in J) (Y-axis – $\log_2$ CPM) in T-ALL samples resulting from thymic transplantation (green) and BM transfer (red), as well as WT thymocytes (black). The bright red triangle represents median expression.

**Supplementary Figure 7. Thymus composition of individual mice.** Thymus grafts of the mice depicted in Fig. 5 were analyzed for the origin of the thymocytes within the CD4CD8 double positive population like shown in Fig 5B. A) Each bar represents one mouse, and the thymocyte origin is depicted by different colors, as shown, corresponding to the indicated percentages. B) Relation between the percentage of CD4CD8 double positive cells in graft-derived thymocytes versus the effective level of engraftment measured in bone marrow LSK. C) Relation between the percentage of graft-derived cells in total CD4CD8 double positive thymocytes versus the percentage of CD4CD8 double positive cells in graft-derived thymocytes.
